# Characterization of Eye and Adnexal Tissues in Dogs and Wolves: A Histological and Lectin-Histochemical Approach

**DOI:** 10.1101/2024.07.26.605385

**Authors:** Abel Diz López, Mateo V. Torres, Fabio Martínez Gómez, Ana López-Beceiro, Luis Fidalgo, Pablo Sanchez-Quinteiro, Irene Ortiz-Leal

## Abstract

This study explores the ocular anatomy and glandular components of domestic dogs compared to their ancestor, the wolf, with the aim of identifying evolutionary changes due to domestication and their implications for ocular pathologies. Utilizing histological and histochemical techniques, including hematoxylin-eosin, PAS, Alcian Blue, and lectins, this research conducts a detailed analysis of the canine and wolf ocular systems, particularly focusing on the eyelids, tarsal glands, and conjunctival tissues. Results indicate significant histological differences between the two species, particularly in the thickness and secretion levels of the conjunctival epithelia and the structure of the tarsal glands. Dogs exhibit a thicker epithelium with greater PAS and Alcian Blue positive secretion, suggesting enhanced ocular protection and lubrication adapted to domestic environments. Conversely, wolves display more concentrated glandular secretions and a predominance of acidic mucopolysaccharides, aligning with their adaptation to natural habitats. This study also highlights the translational value of dogs as models for human ocular diseases, given their anatomical and physiological similarities with humans. Such comparisons are essential as they provide insights that can lead to advancements in medical research and clinical applications, especially in the development of treatments for ocular surface disorders.

## Introduction

The eyelids, including the lower, upper and third eyelids, bulbar and palpebral conjunctiva, lacrimal apparatus and their associated glands are not involved in vision, but have a fundamental role in ocular protection in animals (Ansari and Nadeem 2016). Similarly, the ocular surface, which is formed by the corneal, limbal and conjunctival epithelia along with the tear film, plays a vital role in the defense of the eye (Fine and Yanoff 1979). The hermetic barrier formed by the corneal epithelium, which prevents the entrance of pathogens, is transcendental in this defense. This barrier has a weak point, since being a transparent anatomical element, it lacks blood vessels for its nutrition; however, this disadvantage is compensated by the support provided by the limbal tissue (Crespo-Moral et al. 2020). The limbal tissue; as its name suggests, is a transition to the conjunctiva, a highly vascularized mucosal tissue that provides excellent protection against infections and antigens (Gipson and Argüeso 2003). The conjunctiva performs a crucial protective function due to the presence of mucin-secreting goblet cells. These mucins, along with the secretions from the lacrimal glands and the tarsal glands, also known as Meibomian glands, form the tear film (Watanabe 2002). Alterations in goblet cell function change tear composition, leading to various pathologies. Additionally, the conjunctiva-associated lymphoid tissue (CALT) initiates and regulates immune responses, consisting of B and T lymphocytes, macrophages, and dendritic cells (Knop and Knop 2005; García-Posadas et al. 2016).

Since ancient times, dogs have been recognized for their close relationship with humans, a historical coexistence that continues to grow (Sandøe et al. 2015). This relationship has brought many advantages to both species. However, domestication and the intense artificial selection imposed by humans (Smythe 1975) may be responsible for various pathologies that dogs suffer from today, including ocular problems such as conjunctivitis and dry keratoconjunctivitis (Pedraza Aguirre and Beltrán Bareño 2019), as well as other inflammatory and dryness-related conditions in the ocular system. The significance of this component and its evolution is underscored by the fact that inflammatory diseases such as keratoconjunctivitis sicca (KCS) result from abnormalities in the tear film, specifically in the glandular component (Cabral et al. 2005).

To assess the evolution of the ocular system in dogs, comparing it with that of their ancestor, the wolf, could be a revealing approach. This study aims to characterize the eyeball of the domestic dog and the wolf, focusing on its glandular component, and to compare them in search of possible differences that support the hypothesis of artificial evolution due to domestication. To our knowledge, there is no description of the glandular component of the eyeball of the wolf. Furthermore, there is no comparative description between the eyeball of the dog and wild canids, except for the study conducted on the dog and the crab-eating fox (Lantyer-Araujo et al. 2019), in which only routine histological stains were performed.

Additionally, studying the ocular system of canids, holds significant translational value for human ophthalmology. The difficulty of studying and obtaining human tissues makes research in animals essential, as has been done in the pig (Crespo-Moral et al. 2020), and as will be done in this study in the dog. Preclinical animal studies are instrumental in enhancing our understanding of human diseases and advancing medical knowledge. However, traditional laboratory animals such as rabbits, mice, and rats present limitations due to their distinct ocular surface anatomy, physiology, and tear film dynamics when compared to humans (Sebbag and Mochel 2020). The integration of companion dogs into preclinical pharmacology studies addresses these limitations. Dogs, sharing closer anatomical and physiological similarities with humans, serve as valuable models for investigating ocular surface disorders. By utilizing dogs as preclinical models, researchers can better understand the pathophysiology of ocular diseases, evaluate therapeutic interventions, and ultimately improve treatment strategies for human patients. This approach accelerates the development and application of ophthalmic research, bridging the gap between preclinical studies and clinical practice.

The present study will investigate the eyelids, tarsal glands, palpebral conjunctiva, bulbar, fornix, cornea, ciliary body and lacrimal gland in dog and wolves. Sclera, choroid and retina will also be approached in a more complementary way. To carry out this characterization, a macroscopic and microscopic study will be performed using histological and histochemical techniques: hematoxylin-eosin, PAS, Alcian Blue and lectins. The PAS technique (Schiff’s periodic acid) stains neutral polysaccharides and mucopolysaccharides red, while Alcian blue stains acidic polysaccharides blue, thus differentiating two glandular components. Lectins, a technique using agglutinin proteins, are also used to reveal the glycoconjugates. This technique is based on the presence of glycogen in the glands, so their visualization is of great importance in the characterization of the glandular component (Ortiz-Leal et al. 2022b).

## Methods

The eyeballs and their adnexa used in this study were collected from three adult male mixed-breed dogs, *Canis lupus familiaris*, which were sourced from the Rof Codina University Veterinary Hospital at the Faculty of Veterinary Sciences of Lugo. These dogs had died to various clinical conditions. Additionally, eyeballs were obtained from three adult male *Canis lupus signatus* carcasses, which originated from wildlife rehabilitation centers located in the provinces of Galicia and had died due to fatal accidents. The procurement of wolves and fox samples was conducted in compliance with the necessary authorizations from the Galician Environment, Territory, and Housing Department, under the approval codes EB-009/2020 and EB-007/2021.

### Sample dissection and processing

The sample extraction process involved making an oval incision around the eyelids, preserving them since they contain most of the glandular components, and exposing the orbit. Once the conjunctiva was secured with forceps, it was dissected along the bony orbital rim, creating an access orifice to the orbital cavity. Through this opening, we introduced blunt curved scissors to disinsert the eye muscles and the optic nerve, ultimately extracting the eyeball along with all its annexed structures. Once extracted, the eyeballs were immediately immersed in Bouin’s liquid fixative, which has greater penetration power than formalin, better preserves the tissue, and provides more firmness. After 24 hours, the eyeballs were cut into two halves through a sagittal plane that included the anteroposterior axis of the eye. Then, they were transferred to 70% ethanol. The samples were embedded in paraffin and serially cut into 7-micron-thick sections using a rotating microtome. The sections were collected on gelatinized slides and stored in an oven at 50° Celsius.

### General histological stains

For the realization of this study, we have used the following stains:

#### Hematoxylin-eosin (H-E)

After deparaffination and rehydration the slides were stained with hematoxylin for 2 minutes, then washed for 5 minutes in running water and dipped in eosin for 6 minutes. After an additional 5-minute wash, the slides were dehydrated, cleared in xylene and mounted.

#### PAS

Stain of choice for the detection of neutral polysaccharides. It allows differentiation of glycogen-rich and glycogen-poor areas, as well as appreciation of mucous secretions from glandular components. In addition, it is also suitable for the visualization of cartilage and basal membranes (Bera et al. 2017). The basis of the reaction consists of oxidizing the sample tissues with periodic acid and subsequently applying Schiff’s reagent on the sample, which reacts with the aldehyde groups conferring a purple coloration to the mucous glands. The protocol followed is detailed in depth in Torres et al. (2020).

#### Alcian blue (AB)

Alcian Blue is a basic dye that binds to carbohydrates, giving them a bluish coloration, so it is used to differentiate acid mucopolysaccharides. This is why it is often combined with PAS staining to differentiate the type of mucopolysaccharides present in the serous glands of the tissue. The detailed protocol applied is specified in Ruiz-Rubio et al. (2023).

#### PAS-AA

Staining technique used for the labeling of both glycogen and mucopolysaccharides, obtaining a contrast in the tissue between the acid mucopolysaccharides (stained with Alcian Blue) and the rest of mucopolysaccharides (stained with PAS). The procedure used is thoroughly described in Salazar et al. (2003).

### Lectin histochemical labelling

The histochemical labeling performed in this study involved techniques based on the specific binding of lectins. Lectins, primarily of plant origin, are a type of agglutinin that possess domains capable of recognizing and binding to terminal carbohydrates present in tissues (Plendl and Sinowatz 1998). This binding forms biotinylated macrocomplexes, which are then detected using an ABC complex with peroxidase. Contrary to immunohistochemical techniques, lectins do not have an immune origin and are distributed in various animal and plant tissues where they perform a variety of functions (Devi and Basil-Rose 2018). In this study we have employed two different lectins to characterize the glands present in the adnexa of the eyes.

#### LEA

Lectin corresponding to an agglutinin extracted from tomato (*Lycopersicum esculentum*) that has a high affinity for N-acetylglucosamine, which in turn binds to a specific carbohydrate (Ortiz-Leal et al. 2022a).

#### UEA

Lectin corresponding to agglutinin from gorse (*Ulex europeaus*) that recognizes in priority the terminal L-fucose belonging to glycoproteins and glycolipids (Ortiz-Leal et al. 2024).

The rehydrated slides were initially subjected to a blocking step using 2% bovine serum albumin (BSA) to prevent nonspecific binding, followed by labeling with lectins. For UEA labeling, the samples were incubated for 1 h with unconjugated UEA lectin (L-1060; Vector Labs, Burlingame, CA). Subsequently, a peroxidase-bound anti-UEA antibody (Dako, Denmark), was added and the slides incubated overnight. In the case of LEA labelling, a biotinylated lectin (B-1175-1; Vector Labs) was used and incubated overnight. The next day, the slides were incubated for one and a half hours with an ABC complex (Vectastain, Vector Labs), consisting of peroxidase and an avidin-biotin complex, which binds to the lectin. Visualization of the reaction was achieved using a solution of 0.003% hydrogen peroxide and 0.05% 3,3-diaminobenzidine (DAB) in a 0.2 M Tris-HCl buffer, resulting in a brown-colored deposit. The detailed procedure is explained in Ortiz-Leal et al. (2023).

### Imaging and digital processing

To obtain the digital images showed in this study we used the following equipment and computer programs:

#### Photomicroscope

An Axiophot microscope from Karl Zeiss, equipped with a digital camera, model MRc5 Axiocam, allowed us to photograph large sample areas at high resolution. This is achieved by adjusting a series of parameters such as white balance, exposure, focus, and shading prior to capturing image.

#### Automatic fusion program (PTGui, Rotterdam, The Netherlands)

This software allowed us to merge the photographs obtained with the photomicroscope thanks to the common references (control points) that they present between each one, obtaining a final mosaic of excellent quality of all the photographs of the same sample (in variable number from 6 to 300 photos depending on the sample) resulting in the digital image that we will use (Fig. 1).

**Figure 1:**
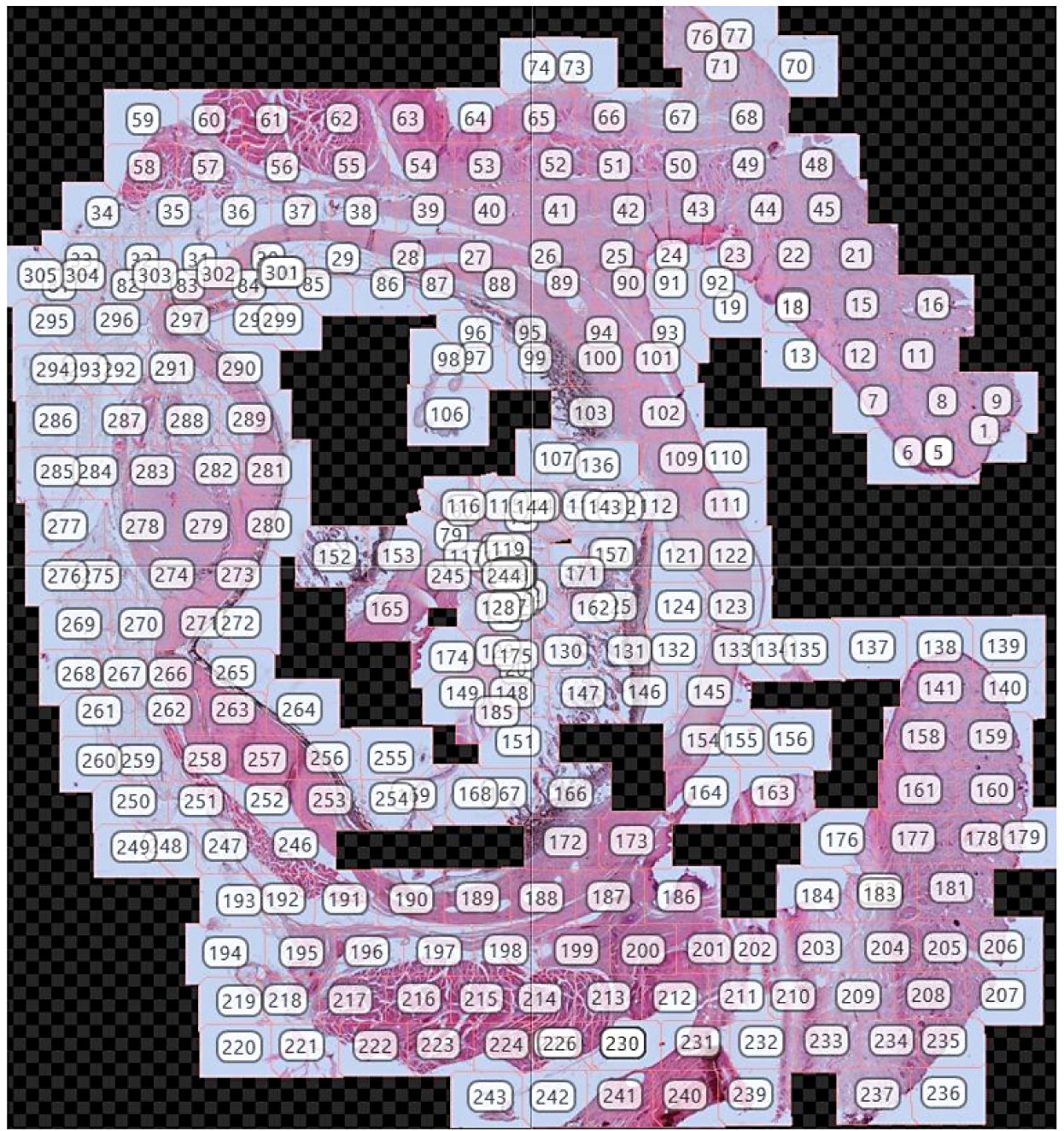
Composite mosaic from 305 photos of a sagittal *Canis lupus* eye’s section.

#### Adobe Photoshop CS4 (Adobe Systems, San Jose, CA)

We used this software for all the images where it was necessary to digitally adjust certain parameters such as contrast, brightness and white balance. No enhancements, additions, or relocations of the image features were made.

## Results

### Macroscopic study

As a preliminary step to the microscopic study, the eyeballs were extracted along with the eyelids and the rest of the ocular adnexa included in the orbit. The high penetrating power of Bouin’s liquid preserved the integrity of the structures within the eyeball without the need to section the orbit. Only after fixation, it was sectioned into two sagittal halves as shown in Figure 2. This image displays the fundamental elements that make up the eyeball: the outer tunic, composed of the sclera and cornea; the middle tunic, composed of the choroid, iris, and ciliary body; and the inner tunic formed by the blind and optic parts of the retina. Moreover, since this fixation method does not alter the shape of the eyeball, the three chambers— anterior, posterior, and vitreous—can be recognized. Macroscopically, no differences were observed between the two species, dog and wolf. All elements accompanying the eyeball were preserved in these specimens. The protective structures around the optic nerve, particularly the Tenon’s capsule or eyeball sheath, were notably difficult to section microtomically, although this did not prevent the completion of full histological sections of the eyeball as shown in the following section (Fig. 3).

**Figure 2:**
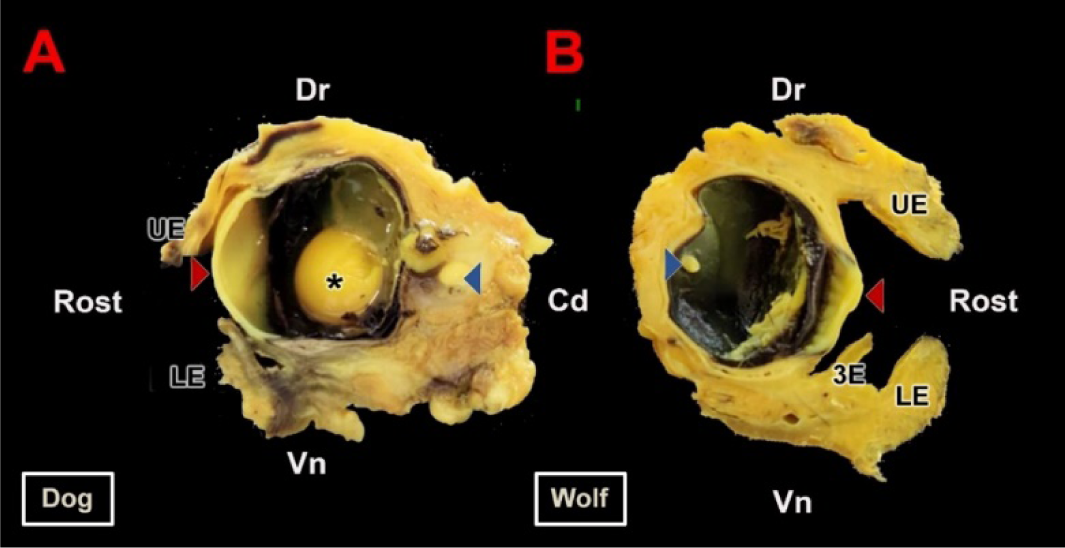
Macroscopic section of the canid eyeball. **A.** Sagittal section of the right eyeball of the dog along its optic axis. **B.** Sagittal section of the left eyeball of the wolf. Samples fixed in Bouin’s fluid and postfixed in 70% ethanol. **Cd**, caudal**; Dr**, dorsal; **LE**, lower eyelid; **UE**, upper eyelid; **Rost**, rostral; **Vn**, ventral; **3E**, third eyelid; **\***, lens; **Red arrow**, cornea; **Blue arrow**, retina.

**Figure 3:**
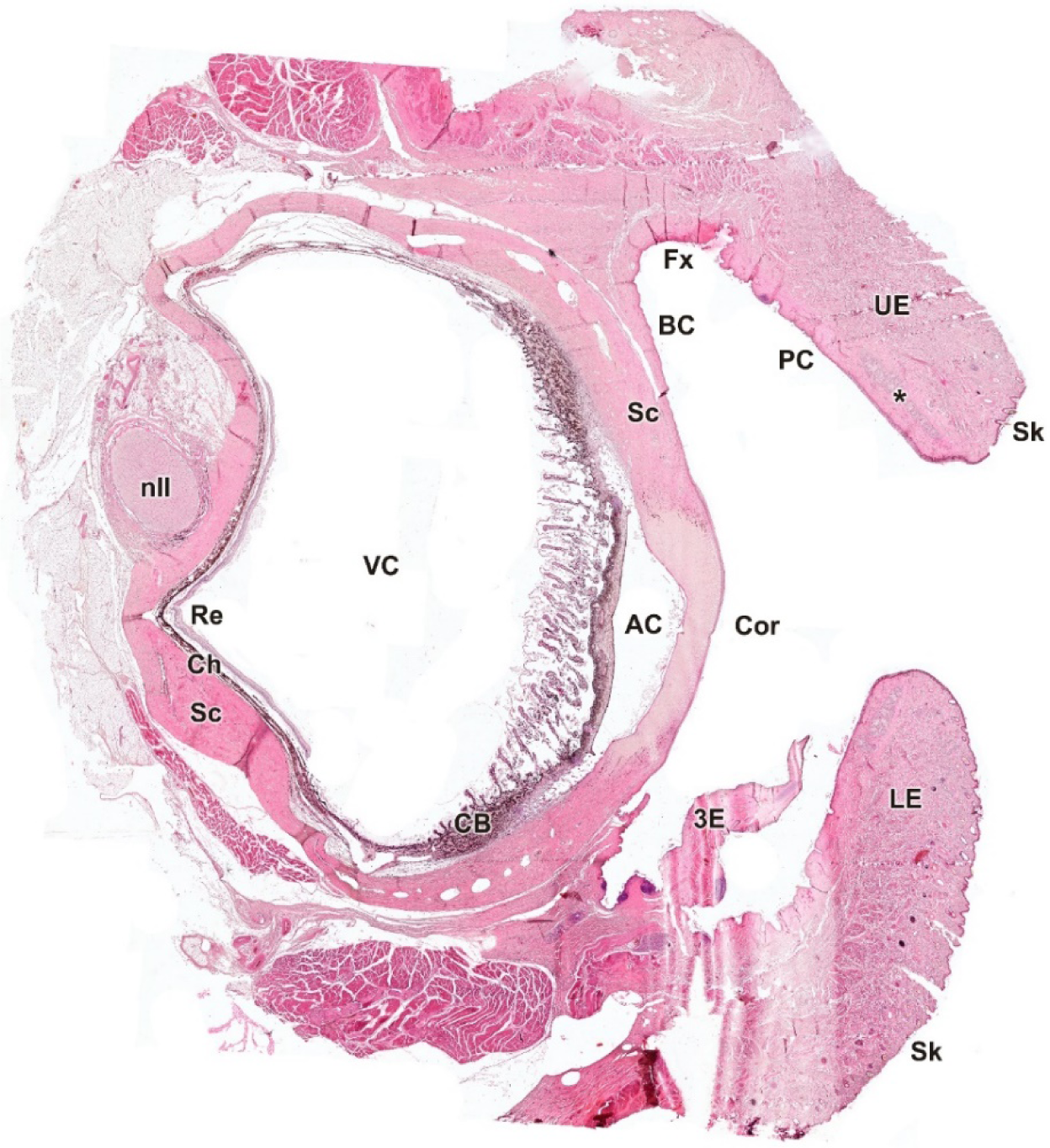
Microscopic sagittal section of the eye of *Canis lupus signatus*. **AC,** anterior chamber**; BC,** bulbar conjunctiva**; CB,** ciliary body**; Ch,** choroid**; Cor**, cornea; **Fx**, fornix; **LE**, lower eyelid; **nII**, optic nerve; **PC**, palpebral conjunctiva; **Re**, retina; **Sc**, sclera; **Sk**, skin; **UE**, upper eyelid; **VC**, vitreous chamber; **3E**, third eyelid; **\***, tarsal glands. H-E staining. Scale bar = 500 µm.

### Microscopic study

As a starting point for the microscopic study, complete sagittal sections of the eyeball and its adnexa were made in the wolf and dog (Fig. 3, 4). The study of these sections permitted to identify the areas responsible for glandular secretion that were the subject of subsequent specific studies with hematoxylin-eosin, Alcian Blue, PAS and PAS-Alcian Blue.

**Figure 4:**
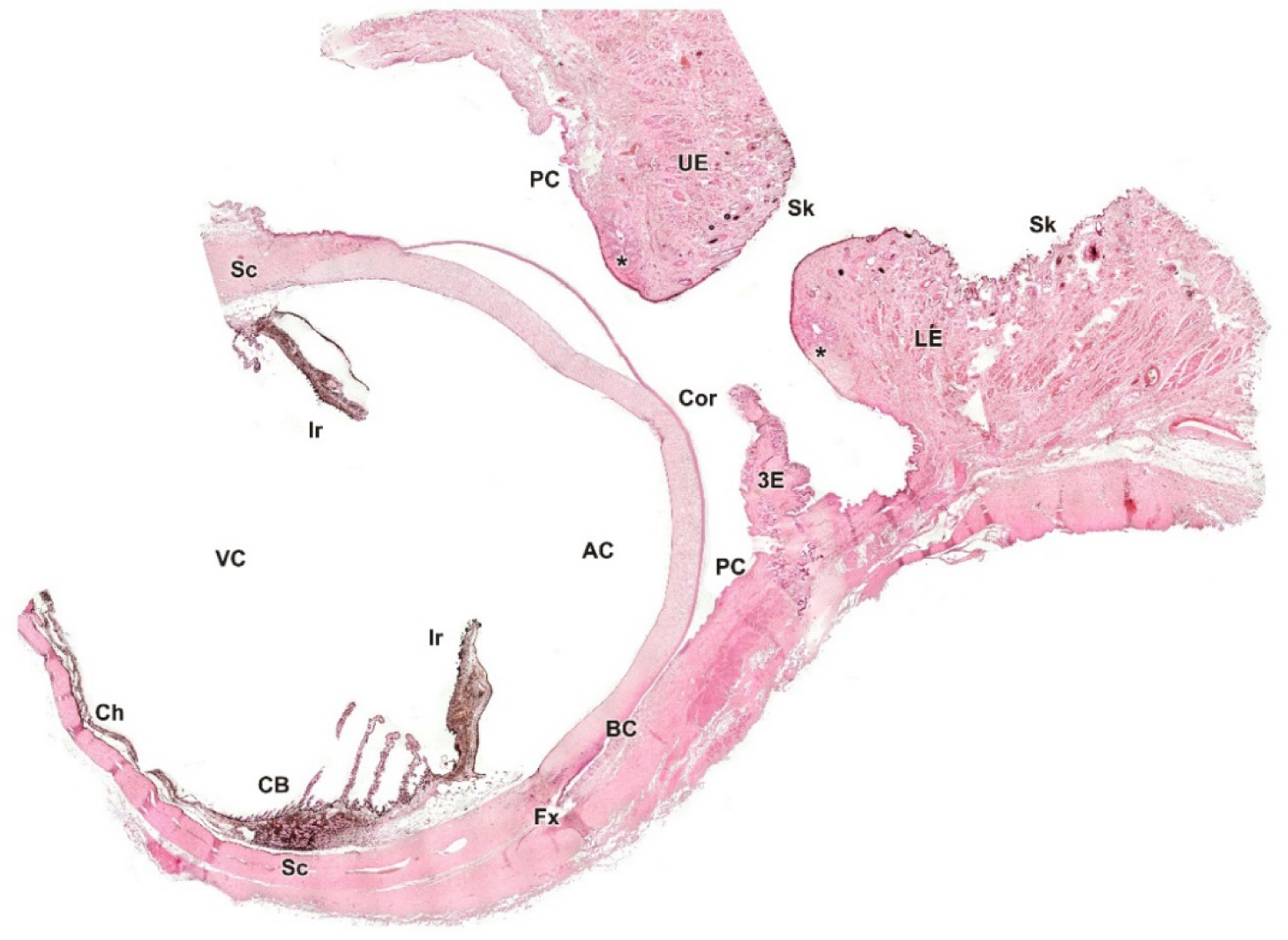
Microscopic sagittal section of the eye of *Canis familiaris*. **AC**, anterior chamber; **BC**, bulbar conjunctiva; **CB**, ciliary body; **Ch**, choroid; **Cor**, cornea; Fx, fornix; **Ir**, iris; **LE**, lower eyelid; **PC**, palpebral conjunctiva; **Sc**, sclera; **Sk**, skin; **UE**, upper eyelid; **VC**, vitreous chamber; **3E**, third eyelid; *, tarsal glands. H-E staining. Scale bar = 500 µm.

### Eyelid skin lining

Upon dissecting the eyeball, the examination of the cutaneous lining of the palpebral region revealed a significant development of its dermal layer, which became more discernible after applying various histological stains, including hematoxylin-eosin., Alcian Blue, PAS, and PAS-Alcian Blue. Comparative analysis, as shown in Figure 5, indicates that both dogs and wolves exhibit similar histological characteristics, including non-keratinized stratified squamous epithelium, hair follicles, and connective tissue. Notably, an abundance of glandular elements in the superficial dermal region is evident, with a higher prevalence noted in the dog (Fig. 5C).

**Figure 5:**
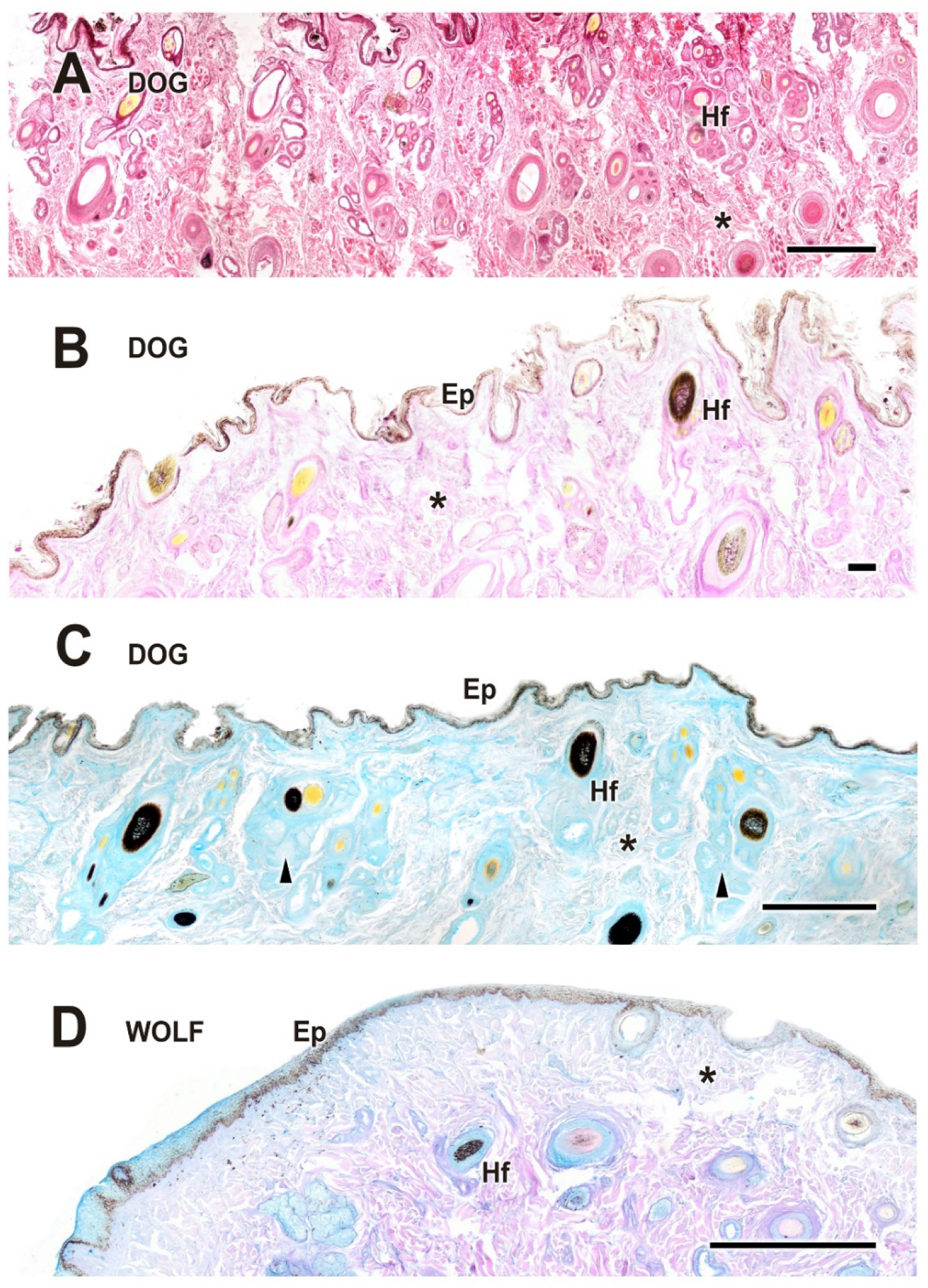
Histological study of the eyelid skin lining in dog (A-C) and wolf (D). The four histological stains used are H-E (**A**), PAS (**B**), AB (**C**), and PAS-AB (**D**), which enable the characterization of the structure of the hair follicles, as well as the epidermis, dermis and connective tissue development. **Ep**, epidermis; **Hf**, hair follicle; arrowheads, tubular glands; *, connective tissue. Scale bar: A, C and D = 500 µm; B = 100 µm.

### Upper eyelid and lower eyelid

The histological examination of the eyelid revealed that both species exhibit fully developed eyelid components, including the epidermis, dermis, hair follicles, and palpebral conjunctiva (Fig. 6). The glandular elements are mainly represented by the tarsal or Meibomian glands, along with the glands of Moll and the glands of Zeis. In both dogs (Figs. 6-8) and wolves (Figs. 7-8), there is a high density of collagen tissue, which is vividly stained with PAS.

**Figure 6:**
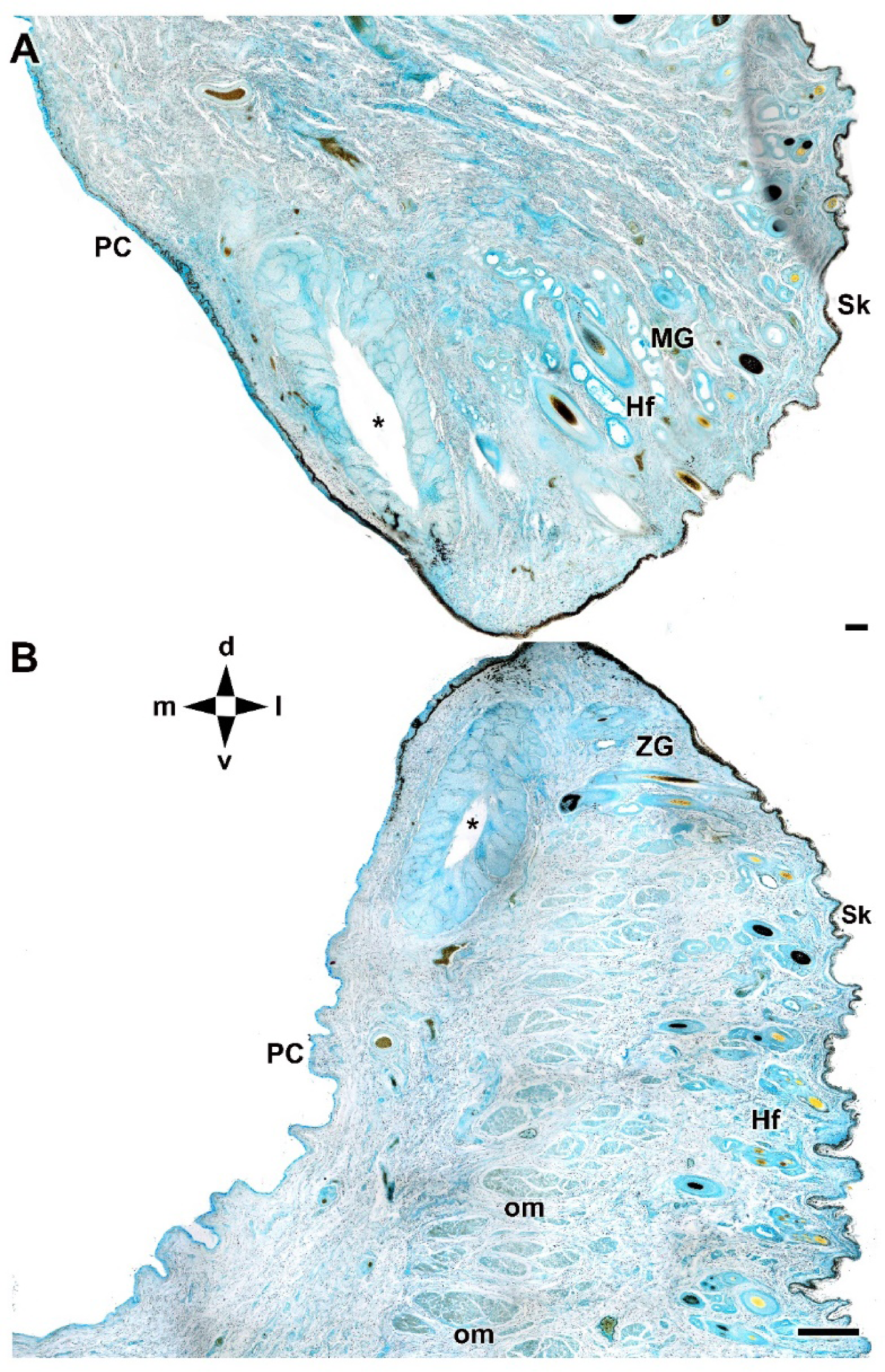
Microscopic anatomy of the dog eyelids. The cross section allows identification of the main glandular elements of the upper (**A**) and lower (**B**) eyelid of the dog. **d**, dorsal; **l**, lateral; **m**, medial; **PC**, palpebral conjunctiva; **Hf**, hair follicle; **MG**, Moll’s gland; **om**, orbicularis muscle; **Sk**, skin; **v**, ventral; **ZG**, Zeis gland; **\***, tarsal glands. AB staining. Scale bar: A = 100 µm; B = 500 µm.

**Figure 7:**
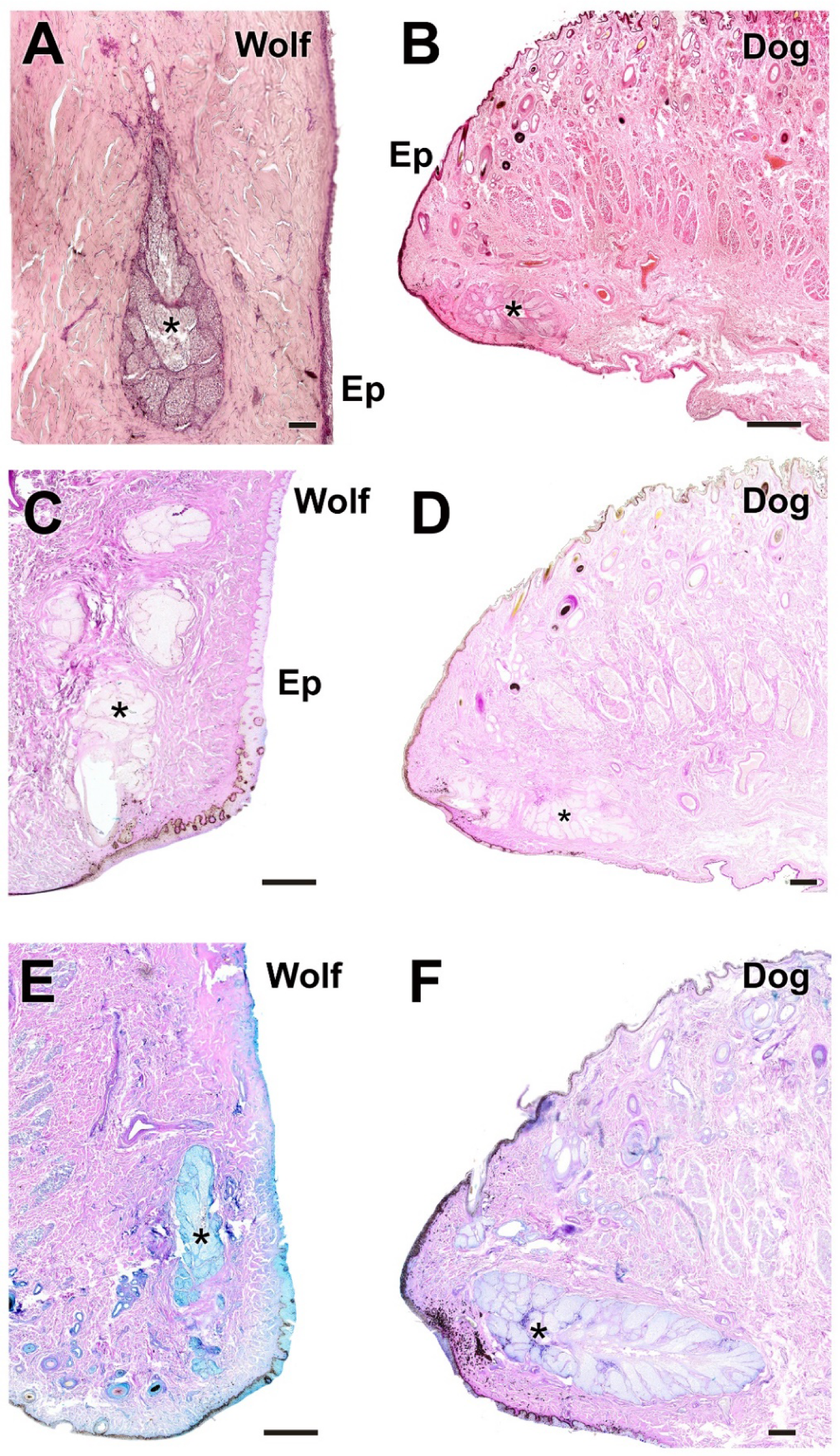
Histological images of dog and wolf upper eyelid stained with H-E, PAS and PAS-AB. They show the main glandular element of the dog (**B**, **D** and **F**) and wolf (**A**, **C** and **E**) upper eyelid, the tarsal glands (**\***). Ep, epidermis. Histological stains: H-E (**A-B**); PAS (**C-D**); PAS-AB (**E-F**). Scale bars: A, D and F = 100 µm; B, C and E = 500 µm.

**Figure 8:**
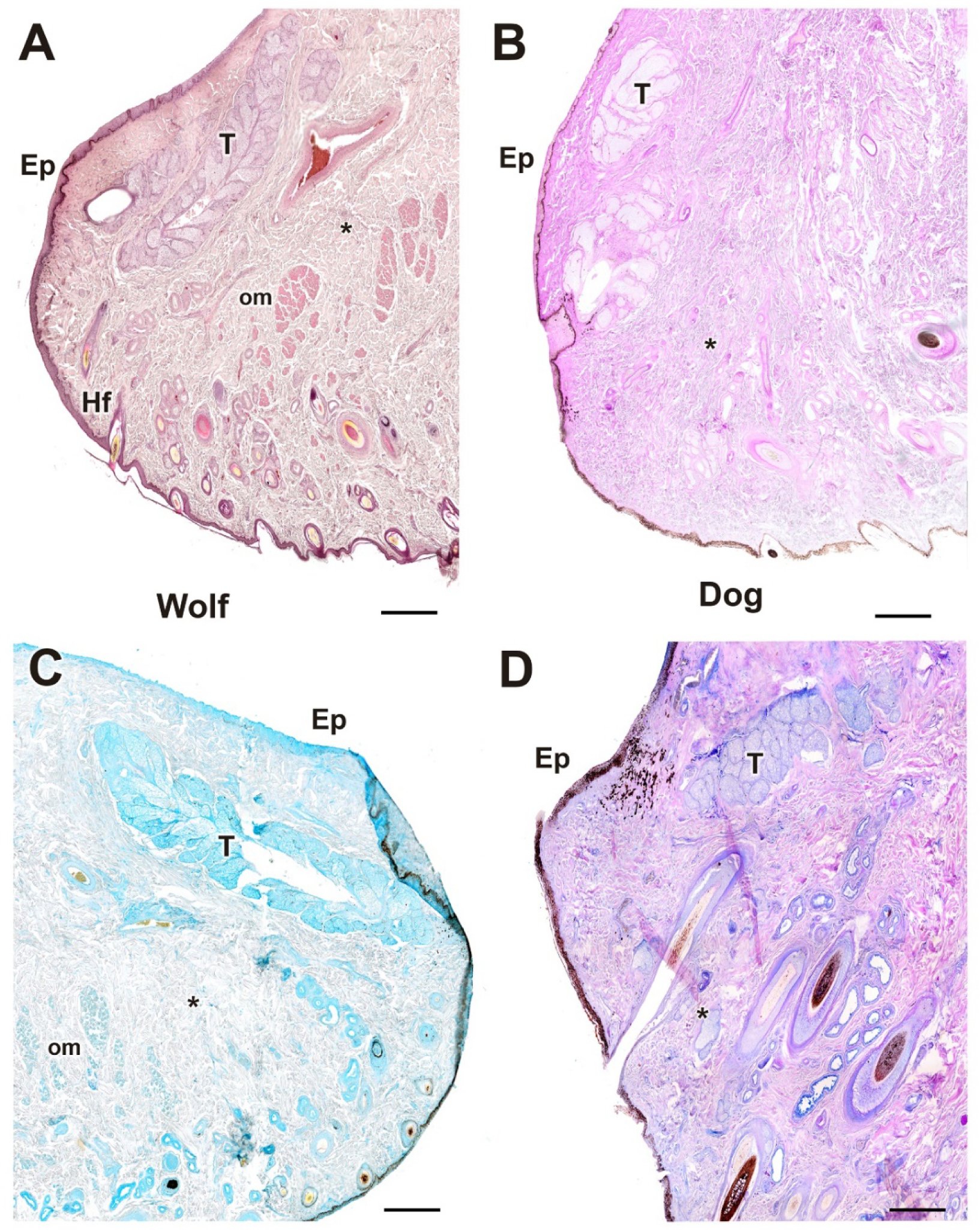
Histological images of dog and wolf lower eyelid stained with H-E, PAS, AB and PAS-AB. The four histological stains used (**A**, H-E; **B**, PAS; **C**, AB; **D**, PAS-AB) allow identification of the main glandular elements of the dog (**B** and **D**) and wolf (**A** and **C**) lower eyelid: **Ep**, epidermis; **T**, tarsal gland; **\***, connective tissue. Scale bar: A-D = 100 µm

This tissue is interspersed with numerous bundles of orbicularis muscle fibers. Remarkably, in dogs, there is a greater development of connective tissue, accounting for the increased elasticity and relative ease of cutting the eyelid compared to wolves, which exhibit a more fibrous character.

### Third eyelid

The histological study of the third eyelid (Fig. 9, 10) has allowed to find remarkable differences between the two species under study, highlighting the different degree of development between the wolf (Fig. 10 A-D) and the wolf (Fig. 10 E-H) The different degree of development between the epidermis of both species can be appreciated. The dog has a thicker epithelium and a greater secretion positivity for PAS and Alcian Blue compared to the wolf. The reaction of the cartilaginous tissue to the four stains (Fig. 9, 10 A, C) is remarkable in both species with no significant differences between them.

**Figure 9:**
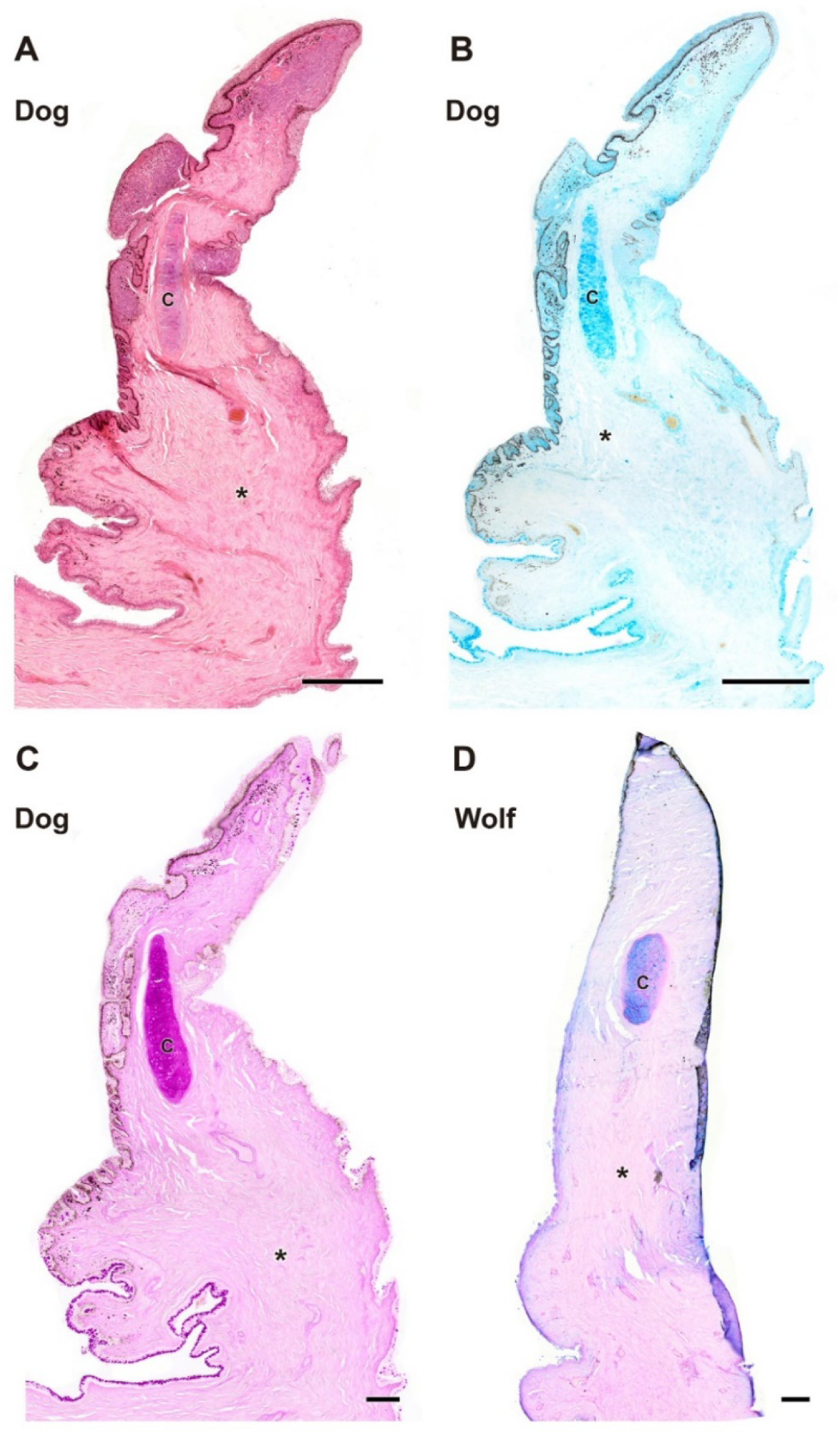
Complete histological sections of the third eyelid of wolf and dog. The stains used (**A**, H-E; **B**, AB; **C**, PAS; **D**, PAS-AB) allow recognition of the cartilaginous skeleton, connective tissue development and epithelial lining epithelia of the bulbar and palpebral surface of the dog (**A**, **B** and **C**) and wolf (**D**): **C**, cartilage; **\***, connective tissue. Scale bar: A and B = 500 µm; C and D = 100 µm.

**Figure 10:**
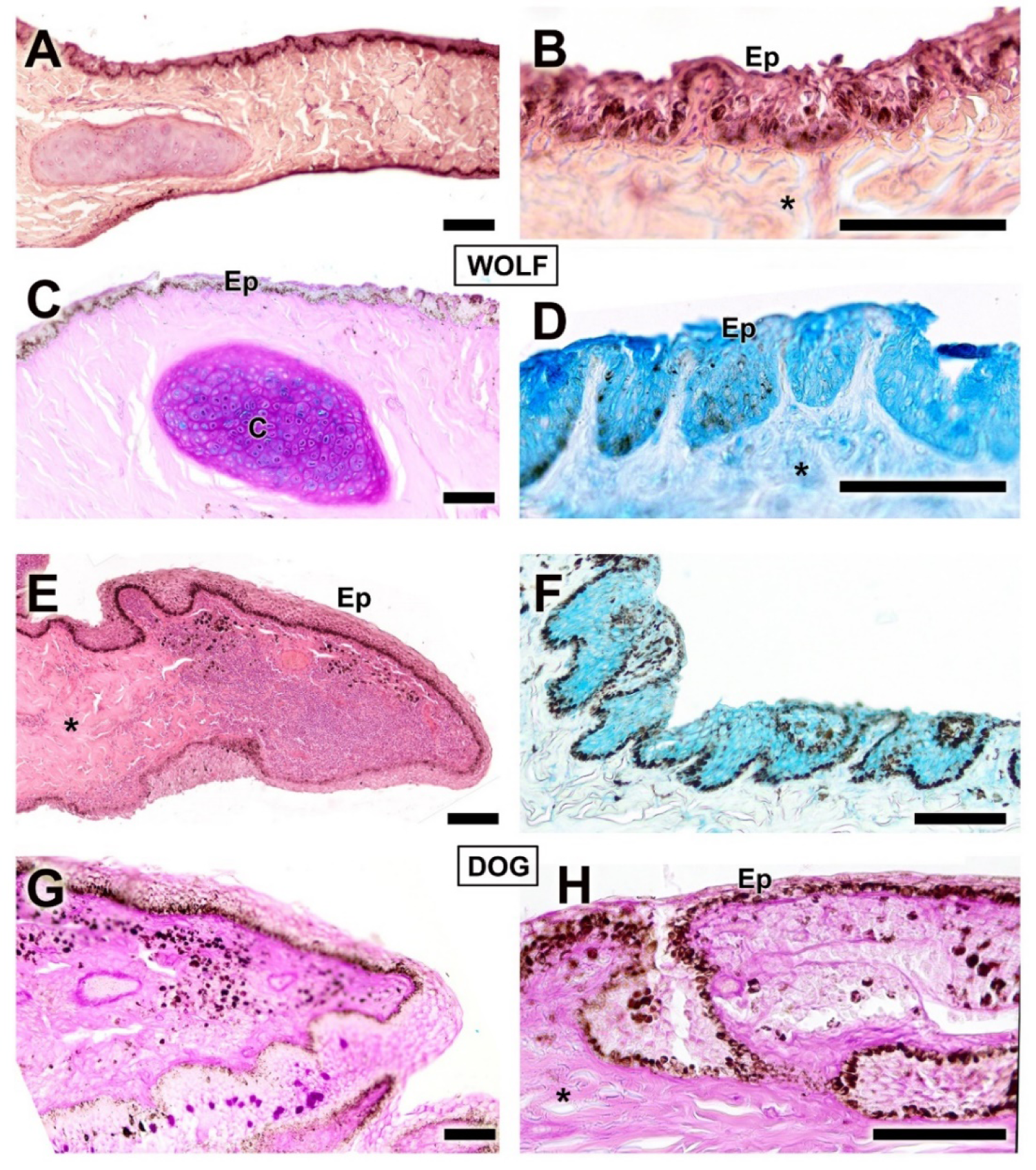
Histological transversal sections of the third eyelid of dog and wolf. The different degree of development between the epidermis of both species can be appreciated. **C**, cartilage; **Ep**, epidermis; **\***, connective tissue. Histological stains used A, B and E, H-E; C, G and H PAS; D and F, AB. Scale bar: A - H = 100 µm.

### Tarsal glands

The tarsal glands in the four stains do not evidence significant differences in the organization of their parenchyma (Fig. 11). However, in dogs, the prominent development of the tarsal lumen is notable (Fig. 11B), while in wolves, the presence of additional aggregates of tarsal glandular tissue is evident (Fig. 11F). In both species, the glandular secretion is acidic in nature and positive for Alcian Blue staining (Fig. 11C and D) but negative for PAS staining, which, in contrast, strongly stains the collagen fibers forming the connective tissue. Remarkably, in wolves, the presence of aggregates consisting of up to three lobes of tarsal glandular tissue is striking (Fig. 11F).

**Figure 11:**
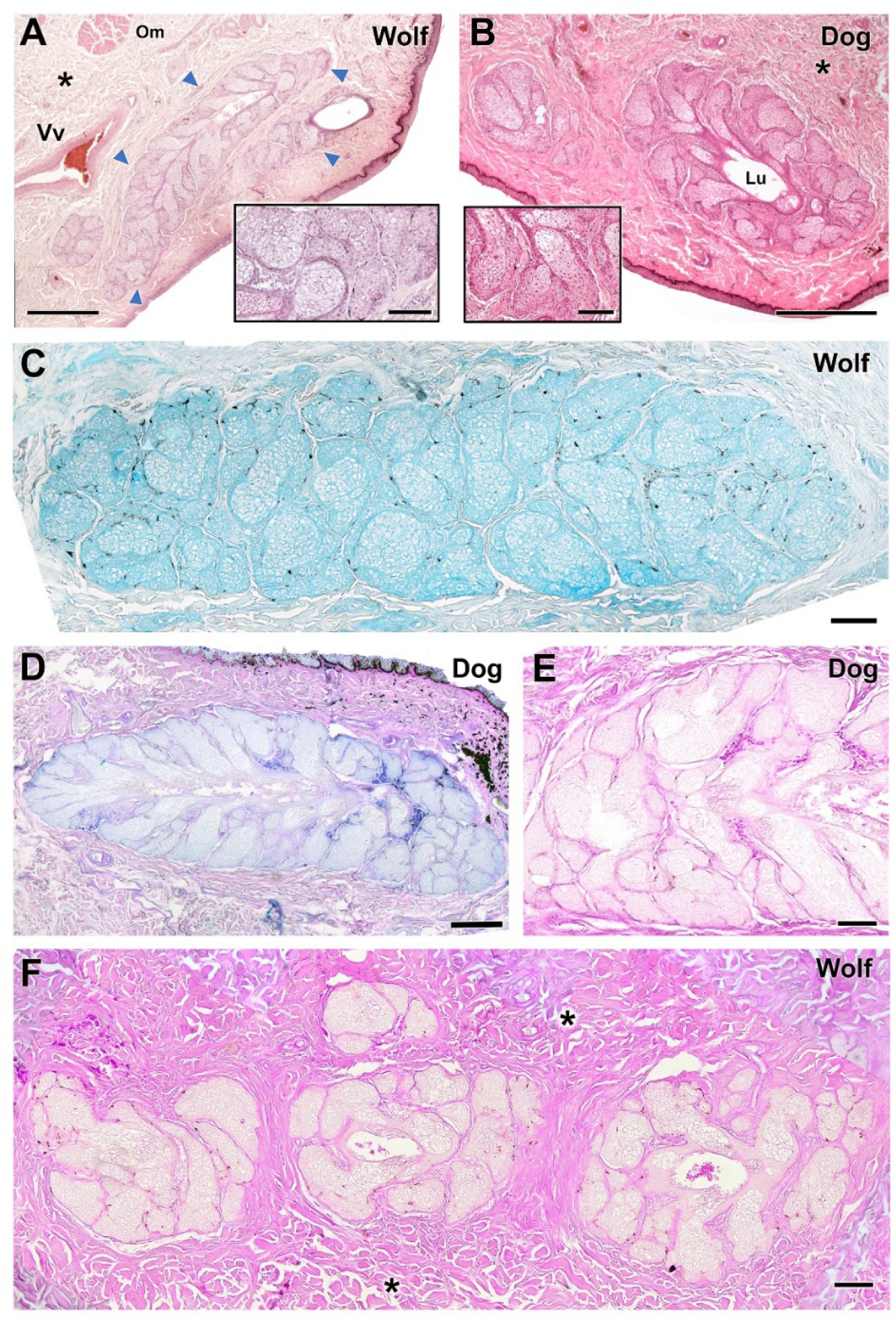
Histological study of tarsal gland of the dog and wolf. **A-B**. H-E staining reveals extensive development of the tarsal gland (arrowheads) in both species with no striking differences. Only in the case of the dog does the large development of the lumen (Lu) stand out. **C-D**. In both species the glandular secretion is acidic in nature, Alcian Blue positive. **E-F**. In both species the tarsal gland is negative to PAS staining which, however, strongly stains the collagen fibers that form the surrounding connective tissue (asterisks). In the case of the wolf, the presence of aggregates of up to three lobes of tarsal glandular tissue is striking. **Om**, Orbital muscle; **Vv**, veins. Staining: A-B, H-E; C, AB; D, PAS-AB; E-F, PAS. Scale bar: A, B, D and E = 500 µm; C and F = 100 µm.

### Palpebral conjunctiva, bulbar conjunctiva and fornix

The histological study of the palpebral conjunctiva, bulbar conjunctiva, and fornix demonstrates intense staining in both types of epithelia in the dog with both Alcian Blue and hematoxylin-eosin. This conjunctiva shows strong pigmentation as well as conspicuous deep tubular invaginations in the palpebral conjunctiva (Fig. 12A, B). In this species, the palpebral mucosa is notably thick (Fig. 13A) and exhibits significant expression of both neutral (Fig. 13B) and acidic mucopolysaccharides (Fig. 13C). In the wolf, acidic mucopolysaccharides predominate in both the palpebral conjunctiva (Fig. 13D) and the fornix (Fig. 13E); this is not the case in the dog, where the fornix primarily expresses neutral mucopolysaccharides (Fig. 13F) in both its connective tissue and unicellular mucous glands.

**Figure 12:**
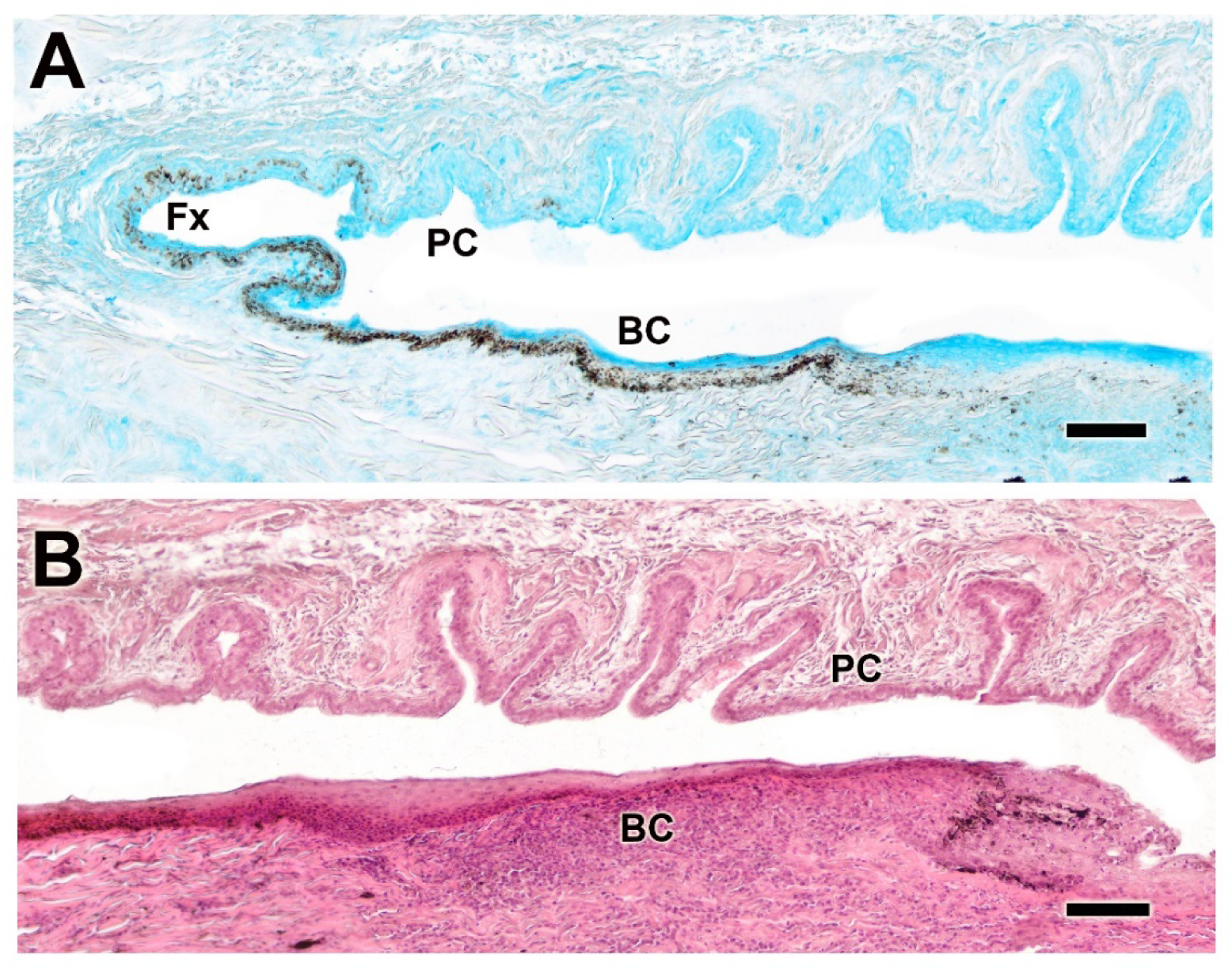
Histological study of the bulbar, palpebral and fornix conjunctivae of the dog. **A**. Alcian Blue staining shows intense staining of the epithelial lining, which is heavily pigmented and forms deep tubular invaginations. **B**. The histological image stained with H-E reveals significant development of these glandular formations, particularly at the level of the palpebral conjunctiva. **BC**. bulbar conjunctiva; **Fx**, fornix; **PC**, palpebral conjunctiva. Scale bar: A and B = 100 µm.

**Figure 13:**
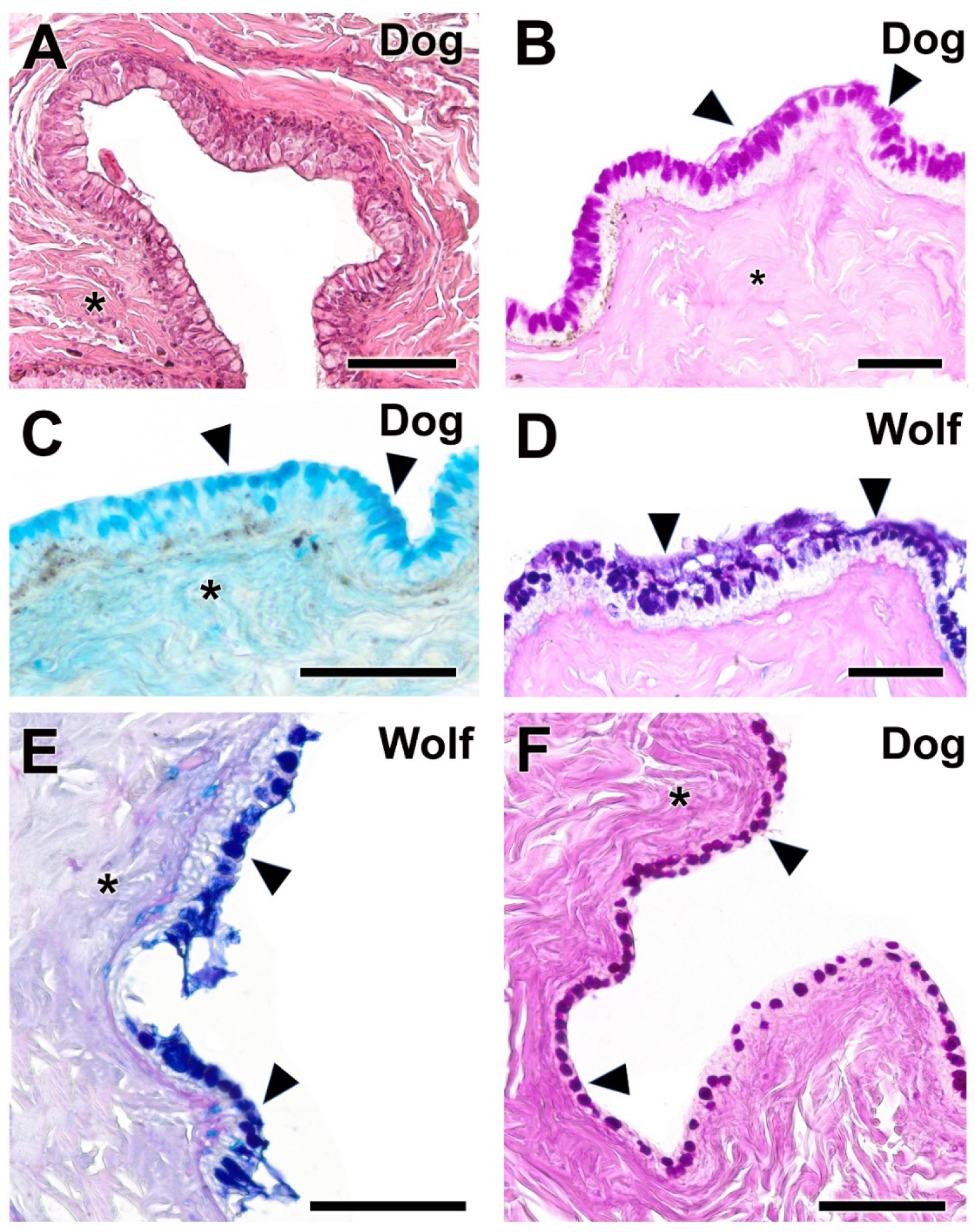
Histological study of palpebral conjunctiva and fornix in dog and wolf. **A**. H-E shows the remarkable thickness of the palpebral mucosa in the dog. **B**-**C**. This morphological development corresponds with a remarkable expression of neutral (**B**) and acidic (**C**) mucopolysaccharides (arrowheads). **D**-**E**. In the wolf, expression of acidic polysaccharides predominates in both the PC (**D**) and the fornix (**E**). **F**. The dog fornix shows a remarkable expression of neutral mucopolysaccharides, both in the unicellular mucous glands and in the submucosal connective tissue (**\***). Stains: **A**. H-E; **B**. PAS; **C**. AB; **D**. PAS-AB; **E**. PAS-AB; **F**. PAS. Scale bar: **A - F** = 100 µm.

The bulbar conjunctiva of the dog (Fig. 14) consists of a highly developed squamous stratified epithelium (Fig. 14A) that shows an intense negative PAS and slight positive AB reaction (Fig. 14C), differing from the wolf. The latter shows a considerably lower thickness and an intense positive AB reaction (Fig. 14B), and slight PAS reactivity mainly on the luminal surface (Fig. 14D).

**Figure 14:**
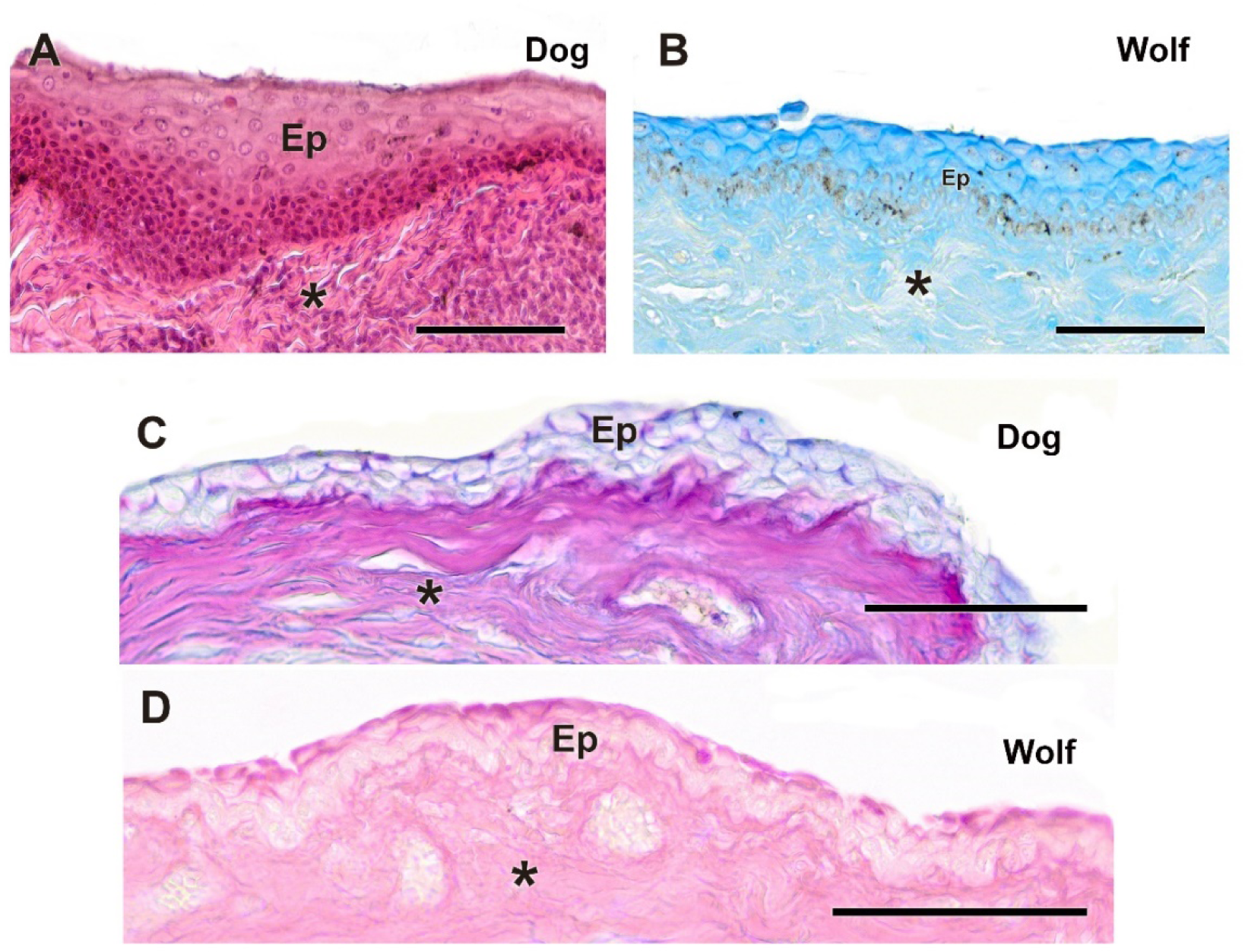
Histological study of the bulbar conjunctiva in dog and wolf. **A**. In the dog, the remarkable development of the stratified squamous epithelium is evident. **B**. In the wolf, the epithelium is thinner and exhibits an intense positive reaction to Alcian Blue. **C**. The bulbar conjunctiva of the dog also shows a strong AB positive reaction and a negative reaction to PAS. **D**. The wolf bulbar conjunctiva shows a certain reactivity to PAS, particularly on its luminal surface. **Ep**. Epidermis; **\***. Connective tissue. Staining: **A**. H-E; **B**. AB; **C**. PAS-AB; **D**. PAS. Scale bar: **A-D** = 100 µm.

### Cornea

The histological study of the cornea does not reveal significant differences between the two species (Fig. 15); however, it allows visualization of the different strata that compose it (Fig. 15B-D). PAS and AB staining show a similar histochemical pattern, with a more intense pigmentation of the stroma in both cases (Fig. 15D, E).

**Figure 15:**
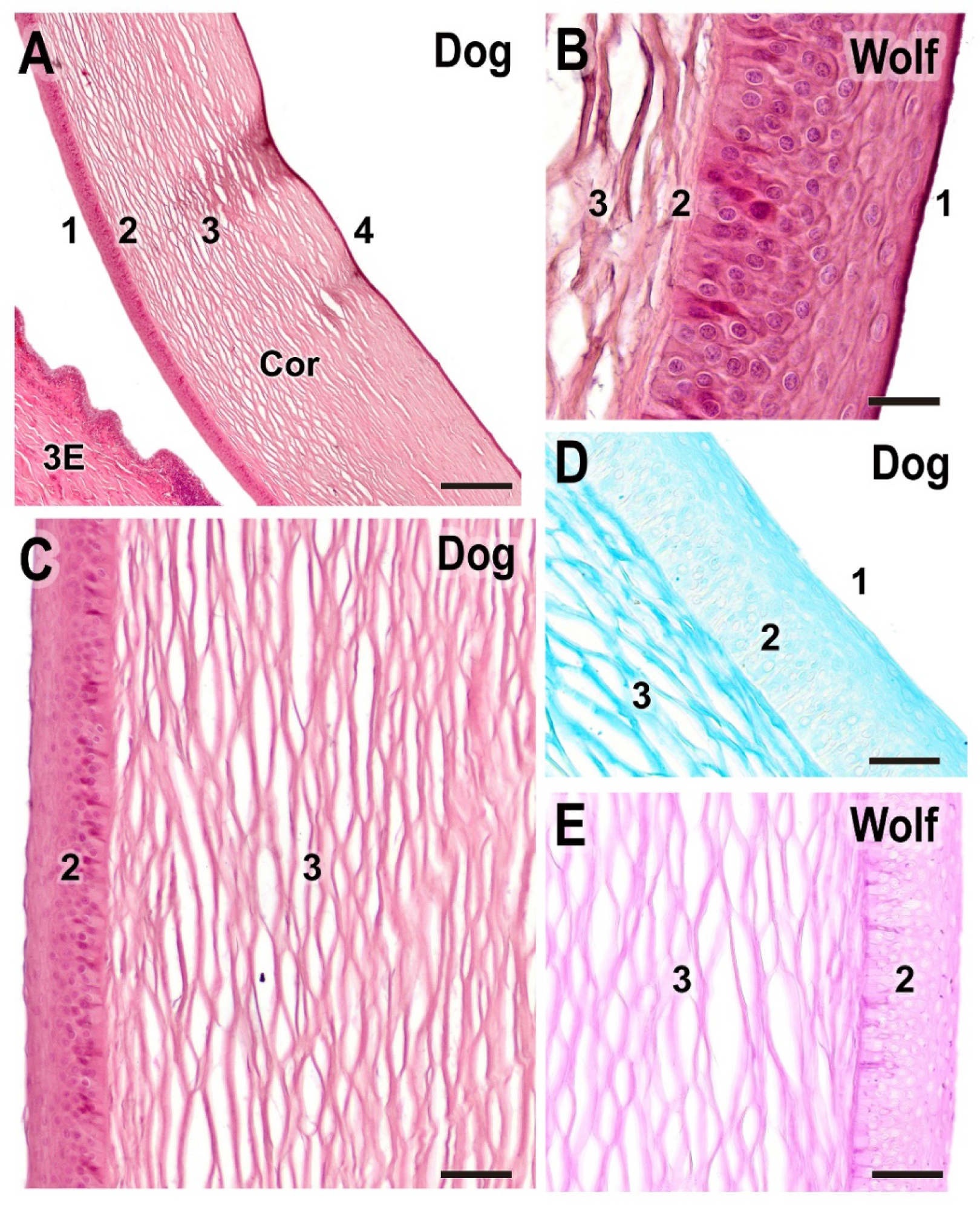
Histological study of the cornea. **A**-**B**. General structure of the dog and wolf cornea, respectively. Sections stained with H-E staining to delineate its various strata. **C**. Histological structure of the superficial corneal strata in the dog stained with H-E. **D**-**E**. Both species show a similar histochemical pattern in this structure, with the stroma staining more intensely in both cases: AB staining in the dog (D) and PAS staining in the wolf (E). **3E**. third eyelid; **Cor**. cornea. 1. anterior epithelium; **2**. anterior limiting lamina (Bowman’s layer); **3**. corneal stroma; **4**. posterior limiting lamina (Descemet’s membrane) + posterior epithelium. **A**-**C**. H-E; **D**. AB; **E**. PAS. Scale bar: **A** = 500 µm; **B**-**E** = 100 µm.

### Ciliary body

The ciliary body is histologically very similar between both species and does not present appreciable morphological differences when stained with Hematoxylin-Eosin, Alcian Blue, and PAS-Alcian Blue, neither for the wolf (Fig. 16A-E) nor for the dog (Fig. 16B-C). However, these stains allow us to observe in detail components such as the ciliary stroma (Fig. 16A), the ciliary processes (Fig. 16A-E), and its two types of epithelia (Fig. 16D-E): a pigmented epithelium with the same degree of pigmentation in both species, and an overlying non-pigmented epithelium, which shows little reactivity to both Alcian Blue and PAS in the wolf (Fig. 16D).

**Figure 16:**
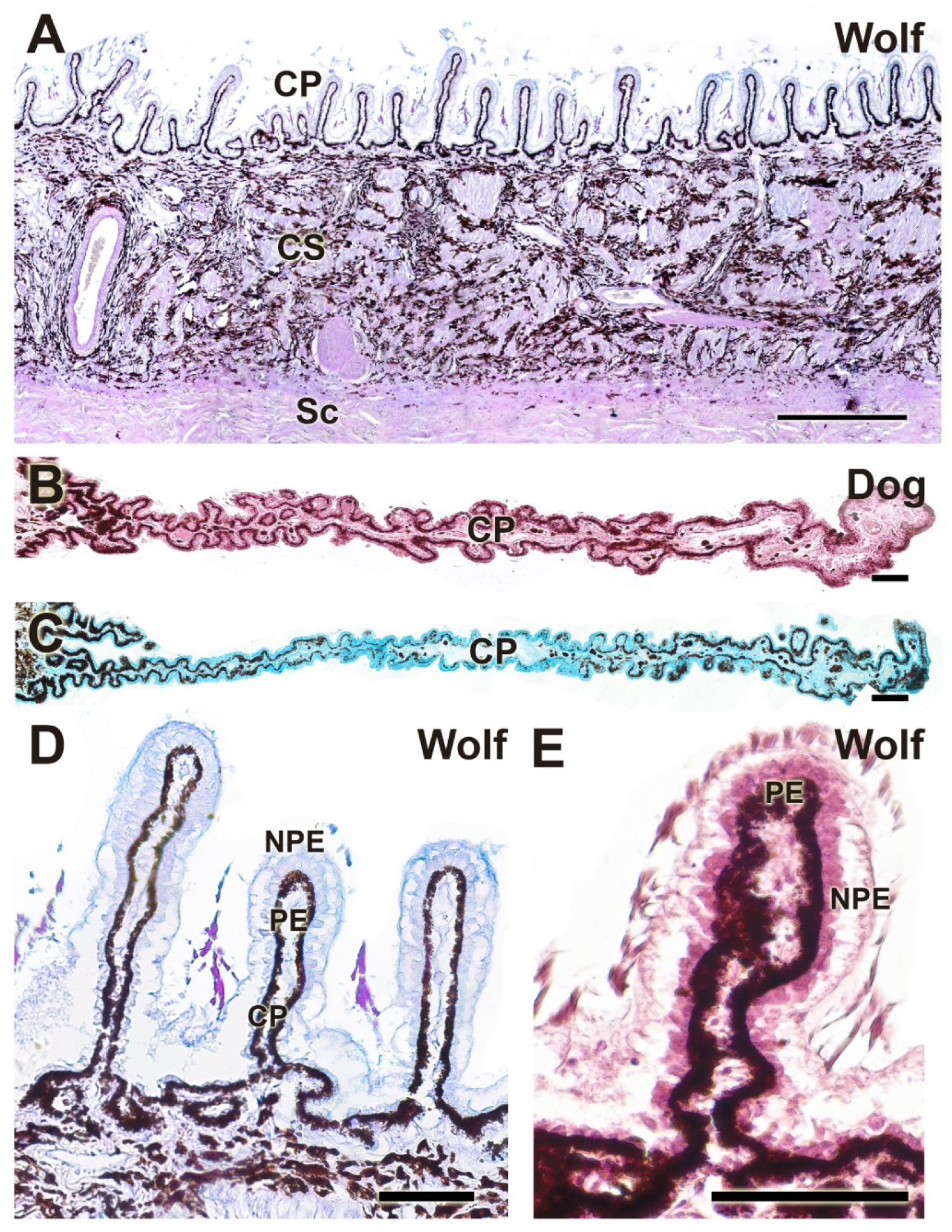
Histological study of the ciliary body. Structurally there are no significant differences between wolf (**A, D, E**) and dog (**B** and **C**). **CP**. ciliary process, **CS**, ciliary stroma; **NP**E, non-pigmented epithelium; **PE**, pigmented epithelium; **Sc**. Sclera. Stains: **A**: PAS-AB; **B, E**: H-E; **C**: AB; **D**: PAS-AB. Scale bar: **A** = 500 µm; **B - E** = 100 µm.

This histological study, conducted using hematoxylin-eosin staining, elucidated the organization of the three principal tunics in the canine eye (Figure 17A), characterized the morphology of the choroid in the wolf (Figure 17B), and delineated the various strata comprising the retina in both species (Figure 17C, D). The retinal layers are organized as follows: 1-Outermost pigmented epithelium; 2-Cone and rod layer; 3-Outer limiting layer, consisting of junctions between photoreceptor cells and Müller cells; 4-Outer nuclear layer, containing nuclei of photoreceptor cells; 5-Outer plexiform layer, a junctional zone between photoreceptor and bipolar cells; 6-Inner nuclear layer, containing nuclei of amacrine, bipolar, and horizontal cells; 7-Inner plexiform layer, a connectivity area among ganglion, amacrine, and bipolar cells; 8-Ganglion cell layer; 9-Optic nerve fiber layer (axons of ganglion cells); 10-Internal limiting layer, primarily serving to separate the retina from the vitreous humor.

**Figure 17:**
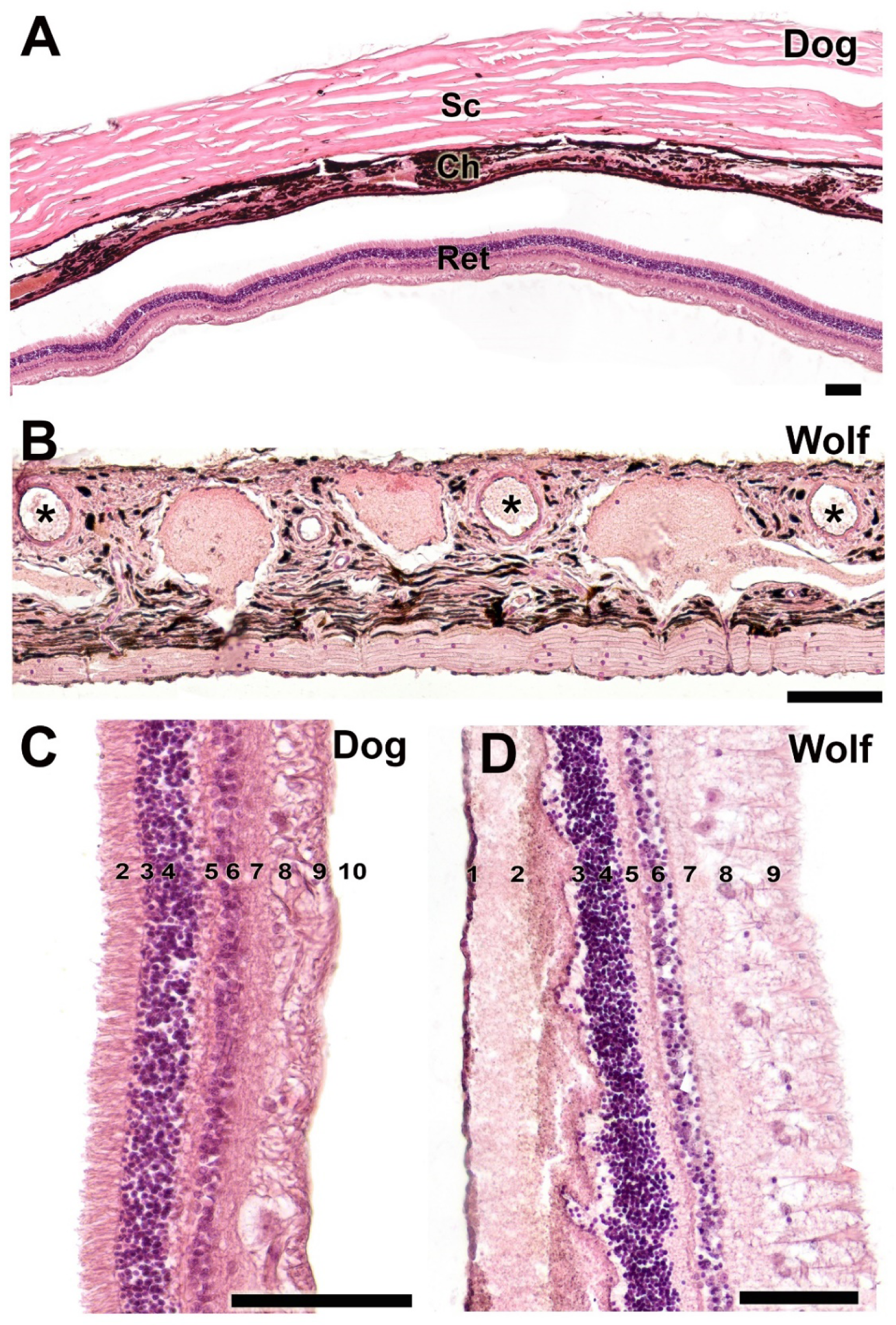
Histological study of the sclera, choroid and retina. **A**. Organization of the three fundamental tunics of the eye in the dog. **B**. Choroid of the wolf. **C**-**D**. Strata that make up the retina of the dog (**C**) and wolf (**D**). **1**. pigmented epithelium; **2**. cone and rod layer; **3**. outer limiting membrane; **4**. outer nuclear membrane; **5**. outer plexiform membrane; **6**. inner nuclear membrane; **7**. inner plexiform membrane; **8**. ganglion cells; **9**. optic nerve fibers; **10**. inner limiting membrane. **Ch**, choroid; **Ret,** retina; **Sc**, sclera; **\***. blood vessels. Staining: H-E. Scale bar: **A - D** = 100 µm.

### Lacrimal gland

In the histological analysis conducted with hematoxylin-eosin staining, we included the morphological organization of the lacrimal gland in the wolf. Special emphasis was placed on the lacrimal duct, which is encased by adipose tissue (Figure 18A), and on the detailed structure of the glandular parenchyma at higher magnification (Figure 18B).

**Figure 18.**
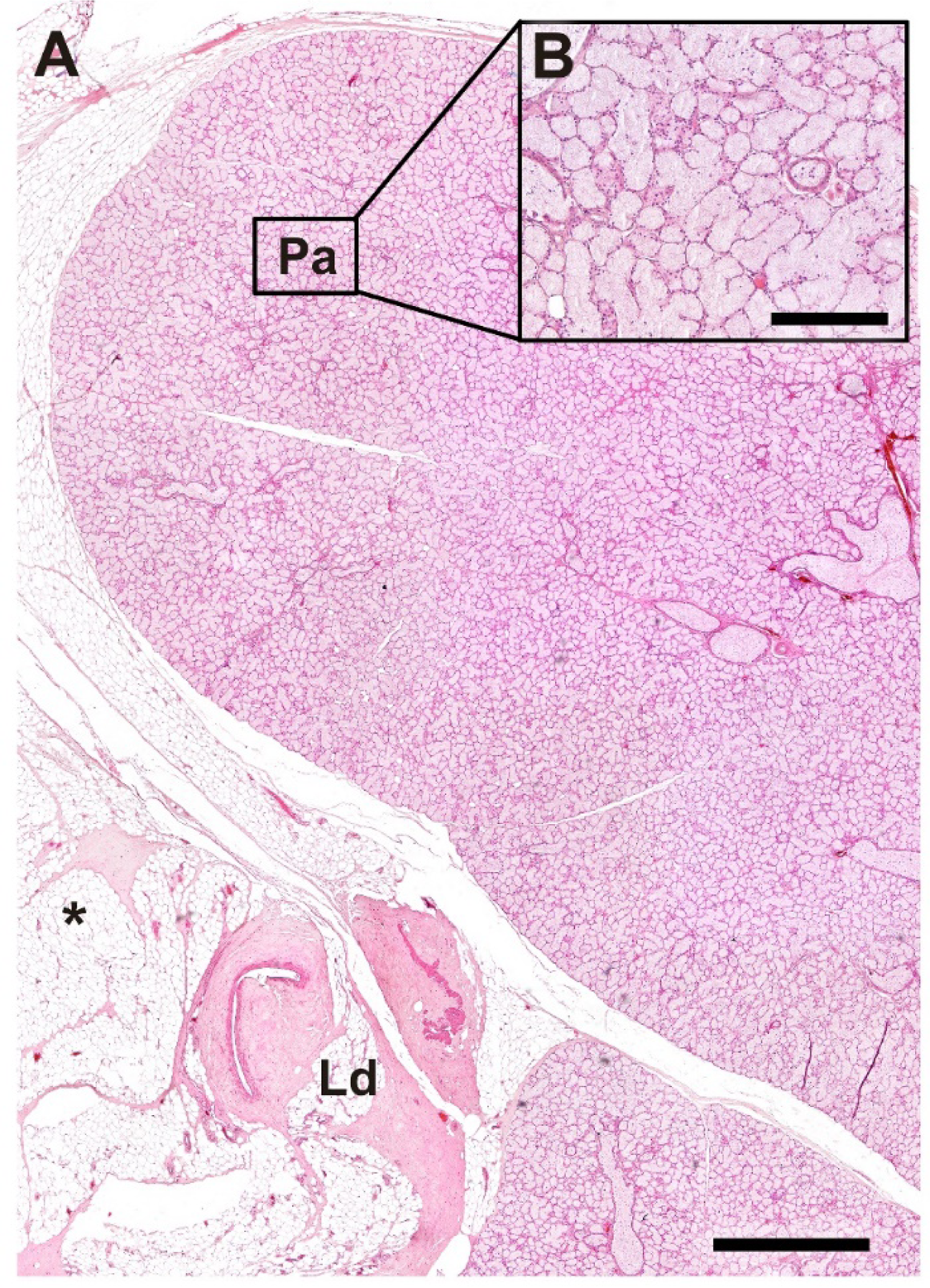
Histological structure of the lacrimal gland of the wolf. **A**. General microscopic view of the gland. **B**. Image of the parenchyma at higher magnification. **Ld**. lacrimal duct; **Pa**. Parenchyma; **\***. Adipose tissue; Stain: H-E. Scale bar: **A** = 500 µm; **B** = 100 µm.

### Lectin histochemical study

The histochemical study of the canine palpebral complex, conducted using UEA lectin (Fig. 19), demonstrated more intense and specific labeling, particularly within the tarsal glands of both the upper and lower eyelids, showing equal intensity in each. Additionally, intense labeling was observed in the mucosal lining epithelia of the upper eyelid, lower eyelid, and third eyelid. Notably, the third eyelid exhibited an intense reaction along its two lining epithelia, characterized by the pronounced presence of pits containing UEA-positive tubular glands (Fig. 20B).

**Figure 19:**
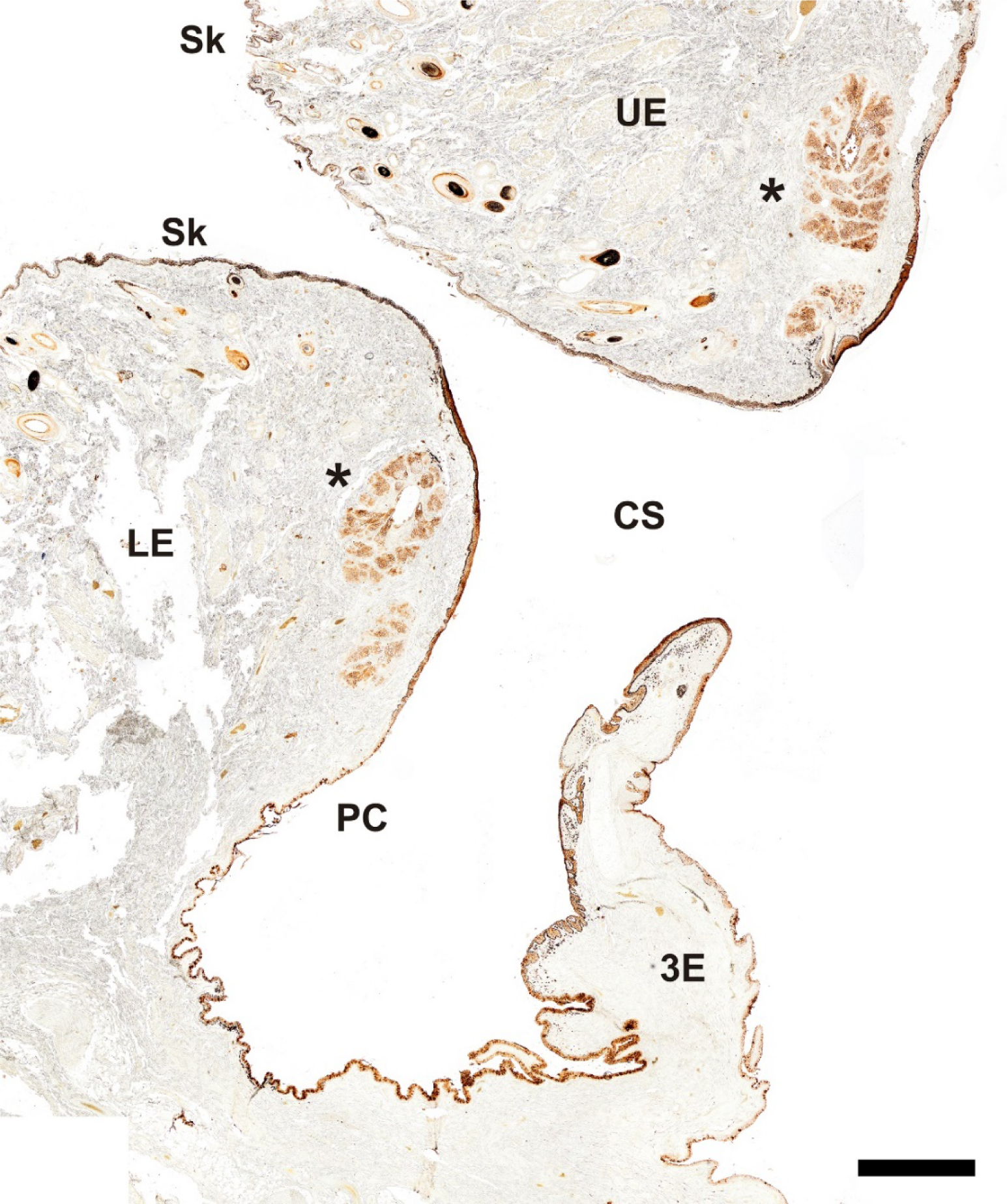
Histochemical study with UEA lectin of the palpebral complex of the dog. The lectin specifically labels the tarsal glands of the upper and lower eyelid, palpebral mucosal linings and hair follicles. **3E**, third eyelid, **CS**, conjunctival sac, **LE**. lower eyelid, **PC**, palpebral conjunctiva; **Sk**, skin; **UE**, upper eyelid. Scale bar = 500 µm.

**Figure 20:**
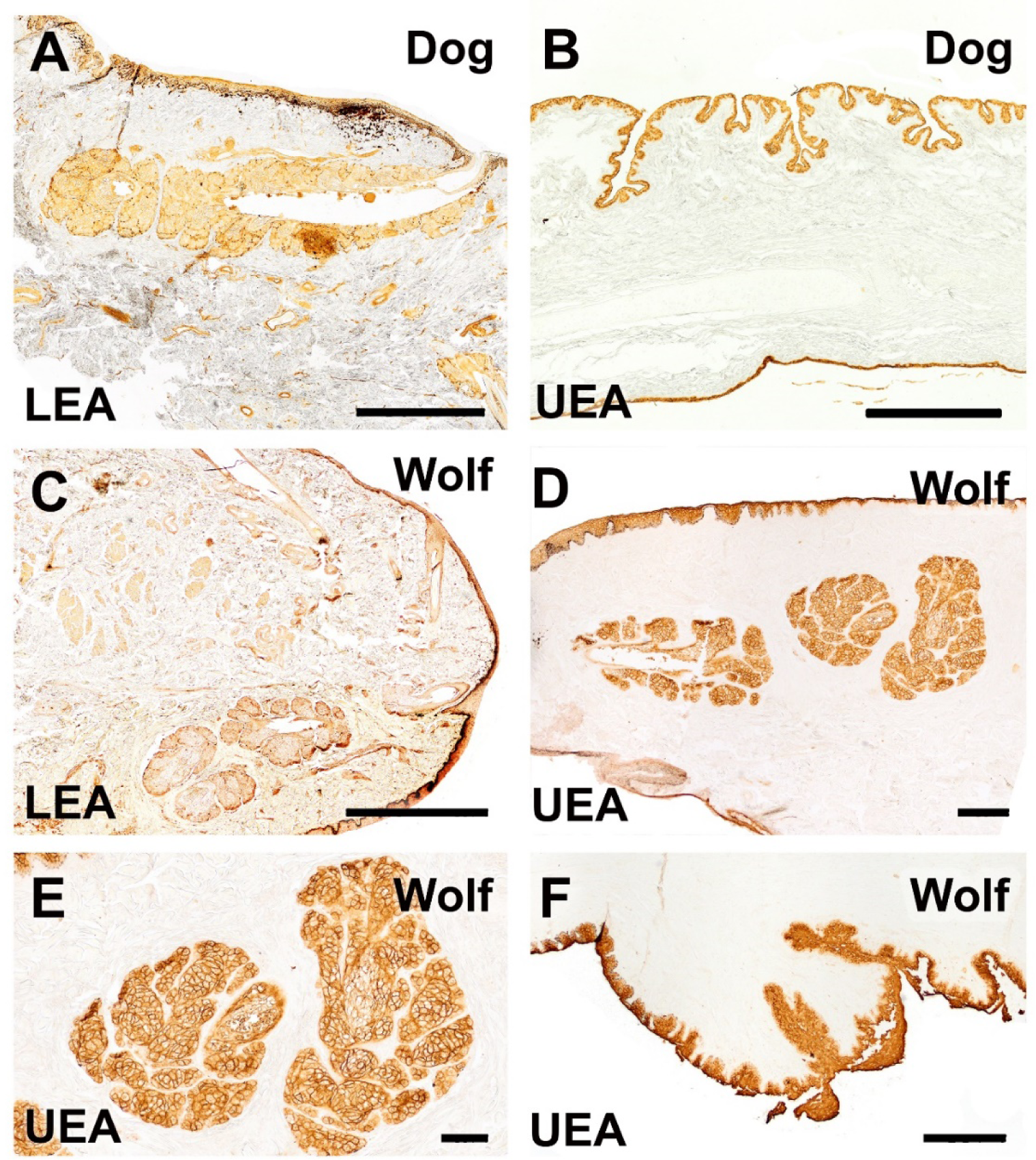
Histochemical study with LEA and UEA lectins of the dog and wolf eyelids. **A** and **C**. LEA lectin exhibits differential labeling between the two species, demonstrating a strongly intense reaction in the tarsal gland of the dog, and a very weak reaction in that of the wolf. **B.** A cross-section of the third eyelid of the dog reveals an intense reaction in its two lining epithelia. Notably, in the palpebral lining, the presence of pits opening into UEA-positive tubular glands is prominent. **D-F**. In the wolf, UEA lectin displays a labelling patern similar to that observed in the dog, with an intense and specific reaction in the skin, palpebral conjunctiva, tarsal glands (shown at higher magnification in E), and the third eyelid (F).Lectins: **A** and **C**. LEA; **B, D, E** and **F**: UEA. Scale bar **A**, **B**, **C** and **F** = 500 µm; **D** and **E** = 100 µm.

Similarly, the UEA lectin labeling pattern in the wolf eye showed no significant differences compared to that observed in the dog, highlighting a specific and strong reaction in the skin, palpebral conjunctiva, tarsal glands, and third eyelid (Fig. 20D-F).

Results obtained with LEA lectin varied between species, presenting differential labeling. A notably intense reaction was observed in the tarsal gland of the canine eye (Fig. 20A), whereas a weaker reaction was noted in the tarsal gland of the wolf eye (Fig. 20C).

## DISCUSSION

In the present study, we have conducted a comprehensive review of the canine eyeball, particularly focusing on the glandular component, which is of vital importance in dogs as these anatomical structures are triggers for common clinical diseases such as keratoconjunctivitis sicca (KCS) or dry eye syndrome (Strom et al. 2016; Montiani-Ferreira et al. 2022). To add a comparative perspective to the study, we have extended it to their ancestor, we extended it to their ancestor, the wolf, attempting to reinforce our hypothesis that artificial selection may have caused significant differences in the ocular anatomy and physiology between wolves and domestic dogs. To this end, we have performed a detailed histological and histochemical description of the ocular apparatus of the wolf, describing for the first time the characteristics of the eyeball and its appendages in this species. The wolf (*Canis lupus*) is listed on the IUCN Red List (IUCN, 2018), underscoring the importance of thoroughly studying and understanding its anatomy and physiology. This knowledge is crucial to lay the groundwork for enhancing the recovery and conservation of this species in its natural habitat. Among the most common ocular diseases in wild canids are conjunctival and corneal pathologies or dry eye syndrome, diseases caused by deficiencies in the aqueous tear component, with many conditions affecting wild canids similar to those observed in domestic dogs (Montiani-Ferreira et al. 2022), which could contribute to worsening the ecological situation of the wolf. This underscores the importance of anatomical knowledge of the eyeball, especially the glandular system focused on in this study.

Our goal is also to provide a translational application, bridging the gap between basic research and clinical practice, contributing to the advancement of research in human ocular pathologies. The domestic canid represents a model much closer to humans than mice and other laboratory animals. Just as other researchers have used the porcine ocular system for similar investigations (Crespo-Moral et al. 2020), the information provided in this study should facilitate research into human ocular pathologies.

Multiple anatomical studies on the ocular system of the dog have been conducted. These studies encompass macroscopic and morphometric characterization of the lacrimal and third eyelid (Cabral et al. 2005), individual analysis of the lacrimal gland (Zwingenberger et al. 2014; El-naseery et al. 2016), evaluation of the cornea and conjunctiva (Strom et al. 2016), the nictitating membrane (Williams and Tighe 2018), and characterization of goblet cells (Moore et al. 1987; Umeda et al. 2010; García-Posadas et al. 2016; Araújo et al. 2019). However, unlike these studies, in our investigation, we have performed an exhaustive analysis combining the study of the ocular surface and eyelids, all supported by a comprehensive and comparative analysis, combining histological and histochemical techniques.

The macroscopic study has confirmed the similarity in both species of the structures comprising the canine eyeball: outer tunic, formed by the sclera and cornea; middle tunic, constituted by the choroid, iris, and ciliary body; and inner tunic formed by the parts of the blind and the optical part of the retina, as well as an examination of the three chambers: anterior, posterior, and vitreous.

Histologically, we have identified the presence of a non-keratinized stratified squamous epithelium, hair follicles, and connective tissue lining the eyelids, consistent with models described by other authors previously (Paszta et al. 2021). Regarding the glandular analysis of the upper and lower eyelids, the presence of the tarsal gland, Moll’s glands, and Zeis glands stands out, also described in the African wild dog, *Lycaon pictus*, a species that could constitute a possible intermediate evolutionary step between the wolf and the domestic dog, as it is a lineage that branched off from the wolf lineage during the Plio-Pleistocene (Hartstone-Rose et al. 2010). In the African wild dog, a stratified squamous epithelium with 9 to 13 layers of nucleated cells is observed, along with elongated tarsian vesicular glands (more developed in the upper eyelids) and the stroma of both eyelids formed by dense and irregular connective tissue; with a network of collagen and elastic fibers (Paszta et al. 2021). This coincides with what we have found in dogs and wolves, where the former species exhibits more developed connective tissue, explaining its greater elasticity compared to that of the wolf, which shows a more fibrous character.

Although both species have a well-developed tarsal gland, the wide lumen in the dog is noteworthy, while in the wolf it may be divided into several independent lobes. The histological study of the palpebral conjunctiva, bulbar conjunctiva, and fornix (reflection of the palpebral conjunctiva onto the bulbar) showed strong pigmentation of both epithelial types in the dog, as well as numerous deep tubular invaginations at the level of the palpebral conjunctiva. This concurs with different studies where, through the quantification of glycoconjugates, it was observed that the highest densities of goblet cells in dogs are found in the palpebral conjunctiva and the fornix (Moore et al. 1987; Gasser et al. 2011). A difference from this study is the low density in the bulbar conjunctiva, which should be less pigmented.

Our histological study has also addressed the organization of the three main tunics in the canine eye: sclera, choroid, and retina, which form the fibrous outer tunic (sclera along with the cornea), the vascular middle tunic (choroid, along with the ciliary body and iris), and the nervous inner tunic (retina; with its optical and blind parts) (Salvador and Martínez 2013).

Lectins selectively bind to cellular surface glycoconjugates, which are involved in cell-to-cell interactions, such as differentiation, regulation, and tissue integrity. The pattern of lectin binding sites exhibits differences not only between the same organs of different species but also within the organs of the same individual. Some of these differences could correlate with various stages of cellular differentiation and organ development (Rittig et al. 1990). The histochemical study of the canine palpebral complex, conducted with UEA lectin, reveals more intense and specific labeling, particularly at the level of the tarsal glands of both the upper and lower eyelids, with equal intensity in both, and at the level of the mucosal lining epithelia of the upper eyelid, lower eyelid, and third eyelid. The latter shows an intense reaction throughout its two lining epithelia, with the notable presence of pits containing UEA-positive tubular glands. The UEA lectin labeling pattern in the wolf’s eye does not show significant differences compared to the labeling observed in the dog, highlighting a specific reaction with a certain degree of intensity in the skin, palpebral conjunctiva, tarsal glands, and third eyelid. The results obtained with LEA lectin differ according to the species, showing differential labeling between them, with a very intense reaction observed in the dog’s eye and a very weak reaction in the wolf’s eye. These studies represent novel findings in both canid species, following the methodologies of various lectin studies conducted in humans (Rittig et al. 1990), bovines (Tuori et al. 1994), and rabbits (Qaddoumi and Lee 2004).

According to various authors (Moore et al. 1987), in both canine and human conjunctiva, glycoconjugates are absent or present in very low amounts on the perilimbal bulbar surface. They are present in greater numbers in the lower nasal fornix, the lower middle fornix, and the lower nasal palpebral area. Conjunctival hydration has been proposed as a significant exogenous factor, with studies suggesting that this degree of hydration is directly related to the number of conjunctival glycoconjugates: This direct relationship is supported by findings from Ralph (1975), which show that the average count of glycoconjugates among patients with dry keratitis was significantly lower than similar counts conducted in normal subjects. These research findings (Ralph 1975; Moore et al. 1987) also support that a high degree of tear fluid hydration is essential for the health of conjunctival goblet cells. Therefore, morphoanatomical studies like the one we have conducted are of critical importance in the fight against clinical ocular pathologies and are an important tool to update and enhance treatments.

At the cutaneous level, both the dog and the wolf display the same non-keratinized stratified squamous epithelium, hair follicles, and connective tissue that covers the eyelids. However, the dog possesses a greater number of glandular components near these areas compared to the wolf. It is notable that in dogs, there is more developed connective tissue, which explains the greater elasticity and ease of cutting the eyelid in this species compared to the wolf, which exhibits a more fibrous character. These findings expand on the evolutionary differences between the two species as suggested by various authors (Montiani-Ferreira et al. 2022). Biologically, this characteristic suggests that dog eyelids are more flexible, facilitating various facial movements and protective functions of the eye. This fact is evident in the great ability of domestic canids to express fear, happiness, or relaxation through the use of facial musculature, as demonstrated by different authors (Caeiro et al. 2017). From a practical perspective in veterinary practice, this elasticity and ease of cutting can simplify surgical interventions, reducing trauma and improving clinical outcomes. Moreover, evolutionarily, this difference may reflect adaptations to domestic environments and specific lifestyles of dogs, in contrast to wolves, which exhibit eyelids with a more fibrous and less elastic character. Various authors (Smith et al. 2024) compare the facial component of humans with that of dogs, showing the facial muscular homology both share, which does not coincide with the wolf, which, as mentioned, exhibits a more fibrous facial component. This is yet another fact that supports the evolutionary differentiation that dogs have undergone due to their shared social environment with humans throughout history

The histological study of the third eyelid has allowed us to identify striking differences between the two species, highlighting the varying degrees of development between the dog and the wolf, both in the lining epithelium of the bulbar conjunctiva and the palpebral conjunctiva. Dogs exhibit a thicker epithelium with greater PAS and Alcian Blue positive secretion than wolves. The tarsal glands show no notable differences in organization; however, in dogs, the extensive development of the tarsal lumen stands out, while in wolves, the presence of additional aggregates of tarsal glandular tissue is noted. In wolves, the expression of acidic mucopolysaccharides predominates in both the palpebral conjunctiva and the fornix, unlike in dogs, whose fornix expresses neutral mucopolysaccharides in both its connective tissue and in the unicellular mucous glands. The bulbar conjunctiva of the dog consists of a highly developed stratified squamous lining epithelium and shows an intense PAS negative and Alcian Blue positive reaction, differing from that of the wolf. In wolves, a considerably thinner epithelium is observed along with an intense Alcian Blue positive reaction that, however, shows some PAS reactivity mainly on the luminal surface. The cornea and ciliary body show no significant differences between the studied species. The sclera, choroid, and retina continue the structural characterization previously described in dogs.

These histological findings reveal significant differences between dogs and wolves with biological, physiological, evolutionary, and veterinary implications. In dogs, the thicker epithelium of the bulbar and palpebral conjunctiva, combined with greater PAS and Alcian Blue positive secretion, may suggest better ocular protection and lubrication capabilities, adapted to domestic environments. The greater development of the tarsal lumen in dogs indicates a greater capacity to store and release secretions. A wider lumen can contain a larger amount of lubricating substances, facilitating their constant and efficient release on the ocular surface. This adaptation allows dog eyelids to maintain better lubrication and ocular protection, especially in domestic environments where they may face various types of irritants and pollutants. On the other hand, wolves, with their additional aggregates of glandular tissue, may have more concentrated and specific secretions, suited for their natural environment, where environmental conditions may require different ocular protection. The predominance of acidic mucopolysaccharides in the palpebral conjunctiva and fornix of wolves, as opposed to the neutral mucopolysaccharides in dogs, reflects adaptations to different environments and ocular protection needs

The histological differences identified are fundamental for the development of specific veterinary treatments and for optimizing the clinical management of ocular conditions in each species. Additionally, these findings provide a solid foundation for future research on ocular adaptations in canids, and are valuable for conservation and species management programs, highlighting the particular adaptations to their respective environments and lifestyles. The potential of this study transcends mere anatomical description, as it offers relevant applications in human medicine. Dogs and humans share many anatomical similarities in the ocular system, as demonstrated by various studies, making the dog an excellent model for surface eye diseases such as dry eye syndrome (KCS) in humans (Strom et al. 2016). This similarity also supports the use of dogs as models for studying the conjunctival mucosal system (Moore et al. 1987). The relevance of these animals in research is not only based on their structural similarities with humans but also on the peculiarities of their evolutionary adaptation, which enhances their utility in the development of pharmaceuticals and ophthalmic devices, as well as in vision science studies. Furthermore, dogs represent the species most commonly affected by spontaneous ocular diseases observed in veterinary practice, serving as significant models for various human ophthalmic pathologies.

In conclusion, this extensive study has not only shed light on the significant histological and evolutionary differences between canines and their wild counterparts but also emphasized the clinical and research potential of these findings. The comparative approach provides critical insights into the anatomical adaptations and health requirements of both domestic dogs and wolves, underscoring the importance of customized veterinary practices and conservation efforts. Furthermore, by highlighting the translational value of dogs as models for human ocular diseases, this research paves the way for future advancements in both veterinary and human medicine. As we continue to explore these relationships, it becomes increasingly clear that understanding the unique characteristics and needs of each species offer opportunities for novel therapeutic strategies and for the improvement of the welfare of both animals and humans.

## AUTHOR CONTRIBUTIONS

Conceptualization, A.D.L., M.V.T., P.S.Q., and I.O.L.; Methodology, M.V.T., F.M.G., P.S.Q., and I.O.L.; Investigation, A.D.L., M.V.T., F.M.G., A.L.B., L.F., P.S.Q., and I.O.L.; Resources, A.L.B., L.F., and P.S.Q.; Writing – Original Draft Preparation, F.M.G., and P.S.Q.; Writing – Review & Editing, A.D.L., P.S.Q, and I.O.L.; Supervision, P.S.Q. and I.O.L.; Project Administration, P.S.Q.; Funding Acquisition, P.S.Q.

## ACKNOWLEDGEMENTS

The authors wish to thank to the wildlife recovery centers from Galicia, and to Dirección Xeral de Patrimonio Natural (Consellería de Medio Ambiente e Ordenación do Territorio, Xunta de Galicia), for having authorized and facilitated the sampling of the animals.

## INSTITUTIONAL REVIEW BOARD STATEMENT

Not applicable, as all the animals employed in this study died by natural causes.

## INFORMED CONSENT STATEMENT

Not applicable, as this research did not involve any humans.

## DATA AVAILABILITY STATEMENT

All relevant data are within the manuscript and are fully available without restriction.

## REFERENCES

Ansari, M.W. and Nadeem, A. 2016. Anatomy of the Eyelids. In: Ansari, M. W. and Nadeem, A. eds. Atlas of Ocular Anatomy. Cham: Springer International Publishing, pp. 53–63. Available at: 10.1007/978-3-319-42781-2_5.

Araújo, R.L.S., Corrêa, J.R. and Galera, P.D. 2019. Ultrastructural morphology of goblet cells of the conjunctiva of dogs. Veterinary Ophthalmology 22(6), pp. 891–897. doi: 10.1111/vop.12667.

Bera, G., Das, R.N., Roy, P., Ghosh, R., Islam, N., Mishra, P.K. and Chaterjee, U. 2017. Utility of PAS and β-catenin staining in histological categorisation and prediction of prognosis of hepatoblastomas. Pediatric Surgery International 33(9), pp. 961–970. doi: 10.1007/s00383-017-4115-2.

Cabral, V.P., Laus, J.L., Dagli, M.L.Z., Pereira, G.T., Talieri, I.C., Monteiro, E.R. and Mamede, F.V. 2005. Canine lacrimal and third eyelid superficial glands’ macroscopic and morphometric characteristics. Ciência Rural 35, pp. 391–397. doi: 10.1590/S0103-84782005000200023.

Caeiro, C., Guo, K. and Mills, D. 2017. Dogs and humans respond to emotionally competent stimuli by producing different facial actions. Scientific Reports 7(1), p. 15525. doi: 10.1038/s41598-017-15091-4.

Crespo-Moral, M., García-Posadas, L., López-García, A. and Diebold, Y. 2020. Histological and immunohistochemical characterization of the porcine ocular surface. PLOS ONE 15(1), p. e0227732. doi: 10.1371/journal.pone.0227732.

Devi, R.V. and Basil-Rose, M.R. 2018. Lectins as Ligands for Directing Nanostructured Systems. Current Drug Delivery 15(4), pp. 448–452. doi: 10.2174/1567201815666180108101246.

El-naseery, N., El-behery, E., El-Ghazali, H. and El-Hady, E. 2016. The structural characterization of the lacrimal gland in the adult dog (Canis familiaris). Benha Veterinary Medical Journal 31(2), pp. 106–116. doi: 10.21608/bvmj.2016.31277.

Fine, B.S. and Yanoff, M. 1979. Ocular histology: A text and atlas. 2nd ed. Medical Dept., Harper & Row.

García-Posadas, L., Contreras-Ruiz, L., Soriano-Romaní, L., Dart, D.A. and Diebold, Y. 2016. Conjunctival Goblet Cell Function: Effect of Contact Lens Wear and Cytokines. Eye & Contact Lens: Science & Clinical Practice 42(2), pp. 83–90. doi: 10.1097/ICL.0000000000000158.

Gasser, K., Fuchs-Baumgartinger, A., Tichy, A. and Nell, B. 2011. Investigations on the conjunctival goblet cells and on the characteristics of glands associated with the eye in the guinea pig. Veterinary Ophthalmology 14(1), pp. 26–40. doi: 10.1111/j.1463-5224.2010.00836.x.

Gipson, I.K. and Argüeso, P. 2003. Role of mucins in the function of the corneal and conjunctival epithelia. International Review of Cytology 231, pp. 1–49. doi: 10.1016/s0074-7696(03)31001-0.

Hartstone-Rose, A., Werdelin, L., De Ruiter, D.J., Berger, L.R. and Churchill, S.E. 2010. The Plio-Pleistocene ancestor of wild dogs, Lycaon sekowei n. sp. Journal of Paleontology 84(2), pp. 299–308. doi: 10.1666/09-124.1.

Knop, E. and Knop, N. 2005. The role of eye-associated lymphoid tissue in corneal immune protection. Journal of Anatomy 206(3), pp. 271–285. doi: 10.1111/j.1469-7580.2005.00394.x.

Lantyer-Araujo, N.L., Silva, D.N., Estrela-Lima, A., Muramoto, C., Libório, F. de A., Silva, É.A. da and Oriá, A.P. 2019. Anatomical, histological and computed tomography comparisons of the eye and adnexa of crab-eating fox (Cerdocyon thous) to domestic dogs. PloS One 14(10), p. e0224245. doi: 10.1371/journal.pone.0224245.

Montiani-Ferreira, F., Moore, B.A. and Ben-Shlomo, G. eds. 2022. Wild and Exotic Animal Ophthalmology: Volume 2: Mammals. Cham: Springer International Publishing. Available at: https://link.springer.com/10.1007/978-3-030-81273-7 [Accessed: 26 July 2024].

Moore, C.P., Wilsman, N.J., Nordheim, E.V., Majors, L.J. and Collier, L.L. 1987. Density and distribution of canine conjunctival goblet cells. Investigative Ophthalmology & Visual Science 28(12), pp. 1925–1932.

Ortiz-Leal, I., Torres, M.V., Barreiro-Vázquez, J., López-Beceiro, A., Fidalgo, L., Shin, T. and Sanchez-Quinteiro, P. 2024. The vomeronasal system of the wolf (*Canis lupus signatus*): The singularities of a wild canid. Journal of Anatomy 00(1–28), p. joa.14024. doi: 10.1111/joa.14024.

Ortiz-Leal, I., Torres, M.V., López-Callejo, L.N., Fidalgo, L.E., López-Beceiro, A. and Sanchez-Quinteiro, P. 2022a. Comparative Neuroanatomical Study of the Main Olfactory Bulb in Domestic and Wild Canids: Dog, Wolf and Red Fox. Animals 12(9), p. 1079. doi: 10.3390/ani12091079.

Ortiz-Leal, I., Torres, M.V., Vargas-Barroso, V., Fidalgo, L.E., López-Beceiro, A.M., Larriva-Sahd, J.A. and Sánchez-Quinteiro, P. 2023. The olfactory limbus of the red fox (Vulpes vulpes). New insights regarding a noncanonical olfactory bulb pathway. Frontiers in Neuroanatomy 16, p. 1097467. doi: 10.3389/fnana.2022.1097467.

Ortiz-Leal, I., Torres, M.V., Villamayor, P.R., Fidalgo, L.E., López-Beceiro, A. and Sanchez-Quinteiro, P. 2022b. Can domestication shape Canidae brain morphology? The accessory olfactory bulb of the red fox as a case in point. Annals of Anatomy -Anatomischer Anzeiger 240, p. 151881. doi: 10.1016/j.aanat.2021.151881.

Paszta, W., Klećkowska-Nawrot, J.E. and Goździewska-Harłajczuk, K. 2021. Anatomical and morphometric evaluation of the orbit, eye tunics, eyelids and orbital glands of the captive females of the South African painted dog (Lycaon pictus pictus Temminck, 1820) (Caniformia: Canidae). Ambrósio, C. E. ed. PLOS ONE 16(4), p. e0249368. doi: 10.1371/journal.pone.0249368.

Pedraza Aguirre, G. and Beltrán Bareño, A.A. 2019. Queratoconjuntivitis seca y cataratas: algunas afecciones oftálmicas comunes en caninos. Available at: https://hdl.handle.net/20.500.12494/8541 [Accessed: 26 July 2024].

Plendl, J. and Sinowatz, F. 1998. Glycobiology of the Olfactory System. Cells Tissues Organs 161(1–4), pp. 234–253. doi: 10.1159/000046461.

Qaddoumi, M. and Lee, V.H.L. 2004. Lectins as Endocytic Ligands: An Assessment of Lectin Binding and Uptake to Rabbit Conjunctival Epithelial Cells. Pharmaceutical Research 21(7), pp. 1160–1166. doi: 10.1023/B:PHAM.0000033002.93967.5f.

Ralph, R.A. 1975. Conjunctival goblet cell density in normal subjects and in dry eye syndromes. Investigative Ophthalmology 14(4), pp. 299–302.

Rittig, M., Brigel, C. and L⍰tjen-Drecoll, E. 1990. Lectin-binding sites in the anterior segment of the human eye. Graefe’s Archive for Clinical and Experimental Ophthalmology 228(6), pp. 528–532. doi: 10.1007/BF00918485.

Ruiz-Rubio, S., Ortiz-Leal, I., Torres, M.V., Somoano, A. and Sanchez-Quinteiro, P. 2023. Do fossorial water voles have a functional vomeronasal organ? A histological and immunohistochemical study. *The Anatomical Record*, p. ar.25374. doi: 10.1002/ar.25374.

Salazar, I., Lombardero, M., Cifuentes, J.M., Quinteiro, P.S. and Aleman, N. 2003. Morphogenesis and growth of the soft tissue and cartilage of the vomeronasal organ in pigs. Journal of Anatomy 202(6), pp. 503–514. doi: 10.1046/j.1469-7580.2003.00183.x.

Salvador, C.R. and Martinez, M.E.G. 2013. Anatomía Veterinaria. 8. Anatomía del ojo (globo del ojo y órganos accesorios) en las especies domésticas. REDUCA 5(2). Available at: https://www.revistareduca.es/index.php/reduca/article/view/1574 [Accessed: 26 July 2024].

Sandøe, P., Palmer, C., Corr, S. and Serpell, J. 2015. History of companion animals and the companion animal sector. John Wiley and Sons., pp. 8–23.

Sebbag, L. and Mochel, J.P. 2020. An eye on the dog as the scientist’s best friend for translational research in ophthalmology: Focus on the ocular surface. Medicinal Research Reviews 40(6), pp. 2566–2604. doi: 10.1002/med.21716.

Smith, H.F., Felix, M.A., Rocco, F.A., Lynch, L.M. and Valdez, D. 2024. Adaptations to sociality in the mimetic and auricular musculature of the African wild dog (Lycaon pictus). *Anatomical Record (Hoboken*, N.J.: 2007). doi: 10.1002/ar.25441.

Smythe, R.H. 1975. The Eye of the Dog. In: Vision in the Animal World. London: Palgrave Macmillan UK, pp. 60–74. Available at: http://link.springer.com/10.1007/978-1-349-02533-6_4 [Accessed: 26 July 2024].

Strom, A.R. et al. 2016. *In vivo* evaluation of the cornea and conjunctiva of the normal laboratory beagle using time- and Fourier-domain optical coherence tomography and ultrasound pachymetry. Veterinary Ophthalmology 19(1), pp. 50–56. doi: 10.1111/vop.12256.

Torres, M.V., Ortiz-Leal, I., Villamayor, P.R., Ferreiro, A., Rois, J.L. and Sanchez-Quinteiro, P. 2020. The vomeronasal system of the newborn capybara: a morphological and immunohistochemical study. Scientific Reports 10(1), p. 13304. doi: 10.1038/s41598-020-69994-w.

Tuori, A., Virtanen, I. and Uusitalo, H. 1994. Lectin binding in the anterior segment of the bovine eye. The Histochemical Journal 26(10), pp. 787–798. doi: 10.1007/BF02388636.

Umeda, Y., Nakamura, S., Fujiki, K., Toshida, H., Saito, A. and Murakami, A. 2010. Distribution of goblet cells and MUC5AC mRNA in the canine nictitating membrane. Experimental Eye Research 91(5), pp. 721–726. doi: 10.1016/j.exer.2010.08.020.

Watanabe, H. 2002. Significance of mucin on the ocular surface. Cornea 21(2 Suppl 1), pp. S17-22. doi: 10.1097/00003226-200203001-00005.

Williams, D.L. and Tighe, A. 2018. Immunohistochemical evaluation of lymphocyte populations in the nictitans glands of normal dogs and dogs with keratoconjunctivitis sicca. Open Veterinary Journal 8(1), p. 47. doi: 10.4314/ovj.v8i1.8.

Zwingenberger, A.L., Park, S.A. and Murphy, C.J. 2014. Computed tomographic imaging characteristics of the normal canine lacrimal glands. BMC Veterinary Research 10(1), p. 116. doi: 10.1186/1746-6148-10-116.

